# DNA O-MAP uncovers the molecular neighborhoods associated with specific genomic loci

**DOI:** 10.1101/2024.07.24.604987

**Authors:** Yuzhen Liu, Christopher D. McGann, Conor P. Herlihy, Mary Krebs, Thomas A. Perkins, Rose Fields, Conor K. Camplisson, David Z. Nwizugbo, Qiaoyi Lin, Nicolas J. Longhi, Chris Hsu, Shayan C. Avanessian, Ashley F. Tsue, Evan E. Kania, David M. Shechner, Brian J. Beliveau, Devin K. Schweppe

## Abstract

The accuracy of crucial nuclear processes such as transcription, replication, and repair, depends on the local composition of chromatin and the regulatory proteins that reside there. Understanding these DNA-protein interactions at the level of specific genomic loci has remained challenging due to technical limitations. Here, we introduce a method termed “DNA O-MAP”, which uses programmable peroxidase-conjugated oligonucleotide probes to biotinylate nearby proteins. We show that DNA O-MAP can be coupled with label-free or sample multiplexed quantitative proteomics, targeted chemical perturbations, and next-generation sequencing to quantify DNA-proximal proteins and DNA-DNA interactions at specific genomic loci. Furthermore, we establish that DNA O-MAP is applicable to both repetitive and unique genomic loci of varying sizes, from kilobase *HOX* gene clusters to megabase alpha-satellite repeats, and that DNA O-MAP can measure proximal molecular effectors in a homolog-specific manner.

## Introduction

Eukaryotic cells store their genetic material in the form of chromatin, a DNA-protein complex. The function of a eukaryotic DNA locus is executed through the cooperation between its nucleotide sequence and the hundreds of protein factors assembled around it. DNA-protein interactions thus play a fundamental role in regulating both the genome’s structure and message storing functions^1^. Therefore, developing methods to decipher DNA-protein interactions in cells has been a focus of technology development efforts for decades^2^. For instance, chromatin immunoprecipitation followed by sequencing (ChIP-seq^3^), which has emerged as a core technology for epigenomics^4^, surveys the genome-wide binding profile of a target DNA-associated protein. ChIP-seq and related technologies (e.g., DamID^5^, CUT&Tag^6^) have produced an abundance of high-quality datasets that enabled the establishment of database consortia such as ENCODE^7,8^ and IHEC^9^, and significantly accelerated chromatin state annotation efforts^10,11^. Such methods, which profile DNA-protein interactions through a protein-centric lens, require the *a priori* knowledge of which protein(s) to target, and rely on the availability of suitable reagents such as antibodies or genetically engineered cell lines. By targeting a single protein at a time, these methods also inherently ignore the context of protein complexes or transient interactions that may be present at a given locus.

In addition to methods that profile the DNA bound by specific proteins, efforts have been dedicated to addressing the inverse problem—identifying the full collection of proteins assembled on a given DNA locus^12–15^. Such methods include the foundational proteomics of isolated chromatin segment (PICh) technology, which uses a biotinylated oligonucleotide (oligo) probe to affinity label specific genomic DNA intervals via *in situ* hybridization (ISH)^16^. To enhance the stability of probe-chromatin interactions throughout the purification workflow, PICh utilizes oligos containing locked nucleic acid residues^17^, which are highly efficient as hybridization probes against repetitive DNA targets but cost-prohibitive to use to target non-repetitive intervals that require dozens to hundreds of probes to properly target a region of interest^18^. As noted in follow-up work, PICh was effective for repeat sequences but would require significant additional work to extend to more complex genomic sequences^19^. Additionally, even with the increased stability gained from the use of locked nucleic acid probes, the probe-chromatin hybrids can be difficult to maintain when coupled with stringent purification washes^19^. As a consequence, an input of one trillion cells was used for a single purification and identification of proteins interacting with telomeres^16^.

To reach a higher degree of enrichment, which is critical for lower abundance DNA targets, an alternative strategy is to directly biotinylate the proteins that occupy a target DNA locus. This biotinylation can be achieved via targeted proximity labeling using promiscuous biotin ligases^20,21^ or the engineered ascorbate peroxidase (APEX/APEX2) enzymes^22,23^. Since the development of APEX, several methods including C-BERST^12^ and GLoPro^13^, have combined APEX with CRISPR genome targeting to endow it with locus specificity. This involves fusing APEX to a catalytically dead RNA-guided nuclease, Cas9 (dCas9) and directing the fusion enzyme to a specific locus of interest by single guide RNAs (sgRNAs). The locus-docked dCas9-APEX biotinylates the neighboring proteins on electrophilic amino acid side chains, such as tyrosine, enabling protein purification and subsequent identification by mass spectrometry. In the case of GLoPro, APEX-based proximity labeling reduced the input required for each replicate analysis to ∼300 million cells—a 10-fold reduction in cell input compared to PICh. Most recently, an approach termed TurboCas^24^ introduced the combination of dCas9 fused to the miniTurbo^25^ biotin ligase, enabling detection of locus-specific proteins from 50 million cells per replicate. Nevertheless, a notable limitation of CRISPR-guided proximity labeling is the requirement of the fusion dCas9-APEX or dCas9-miniTurbo enzyme and sgRNAs in a suitable host cell line. Since a successful locus purification canonically requires tens to hundreds of millions of cells, if not more, most current methods aim to create stable cell lines for this purpose. These requirements limit the use of previous locus proteomics methods since efficient and well-tolerated gene delivery remains a major challenge and considerable effort in primary cells^26^. In addition, the labeling reagents necessary for APEX-based proximity labeling—hydrogen peroxide and biotin phenoxyl radicals— are toxic to cells and living organisms, limiting the use of CRISPR-based peroxidase labeling to cell lines amenable to genetic engineering. Thus, an unmet need exists for extensible methods capable of scaling and profiling multiple genomic loci.

We address these technical limitations by introducing DNA O-MAP, a locus purification method that uses oligo-based ISH probes to recruit peroxidase activity to specific DNA intervals. DNA O-MAP builds on our previously introduced RNA O-MAP^27^ and pSABER^28^ techniques, which target peroxidase activity to specific RNAs and RNA/DNA intervals for purification or visualization, respectively. Here, we describe a cost-effective and scalable bulk hybridization and biotinylation workflows capable of processing millions of cells in parallel in just a few days, and demonstrate that the recovered material is compatible with sample multiplexed proteomics^29^ and drug perturbation. We benchmark our approach by recovering telomere-specific DNA binding proteins after targeting telomeric DNA. We further showcase the scalability of DNA O-MAP by distinguishing the DNA-associated proteomes around human pericentromeric alpha-satellite repeats, telomeres, and mitochondrial genomes in quadruplicates using tandem mass tags^29^. We go on to demonstrate that DNA O-MAP can capture functionally relevant DNA-DNA interactions, read out by DNA sequencing, from 20 kilobase intervals. Additionally, we show that DNA O-MAP can measure the local proteome of non-repetitive elements such as the *HOXA* and *HOXB* gene clusters as well as differential proteomes at these gene clusters before and after chemical inhibition of chromatin regulation. Finally, we show that DNA O-MAP can be applied to discern homolog-specific local proteomes of the active and inactive X-chromosome. We anticipate that the flexible targeting, scalable protocol, and robust labeling capabilities provided by DNA O-MAP will lead to its adoption as a platform technology for uncovering locus-proximal chromatin proteomes.

## Results

### Design of DNA O-MAP

DNA O-MAP is a molecular profiling methodology that combines the targeting flexibility of oligo-based ISH with the ability of horseradish peroxidase (HRP) to catalyze the localized deposition of small biomolecules at sites where it is bound. DNA O-MAP works by recruiting a ‘secondary’ HRP-conjugated oligo to sites where the primary ISH probes are bound. HRP-mediated deposition of biotin at targeted loci enables the pull-down and purification of proximal and chromatin associated proteins and DNA from *trans*-interacting genomic loci. DNA O-MAP reports on both direct DNA-protein interactions and proteins in spatial proximity to a target locus and allows for identification and quantitative comparison of proximal proteins between target genomic loci. As in RNA O-MAP^27^, the specificity of ISH and/or biotinylation can be assessed by microscopy using a small sample of cells immobilized on solid support before the cells enter affinity purification. Importantly, the HRP-conjugated oligo is available via several commercial sources, allowing researchers without the expertise to perform their own conjugations to utilize DNA O-MAP.

### Establishing a scalable in-solution hybridization-biotinylation workflow for DNA O-MAP

During the development of DNA O-MAP, we refined an in-solution hybridization workflow on cells in suspension for cost-efficient genomic labeling in parallel with an in-dish workflow used for RNA O-MAP^27^ (**Figure 1A**). We began with adherent cells grown on multi-layer flasks, each yielding 90-120 million cells, and subsequently released and fixed (4% PFA) in order to be compatible with DNA ISH. Samples can be processed in parallel, thereby increasing the number of samples that could be handled at once. We note that the in-solution version of the protocol reduces reagent costs by ∼1,000-fold relative to conventional ISH protocols performed on solid substrates to further enhance the scalability of DNA O-MAP.

**Figure 1.**
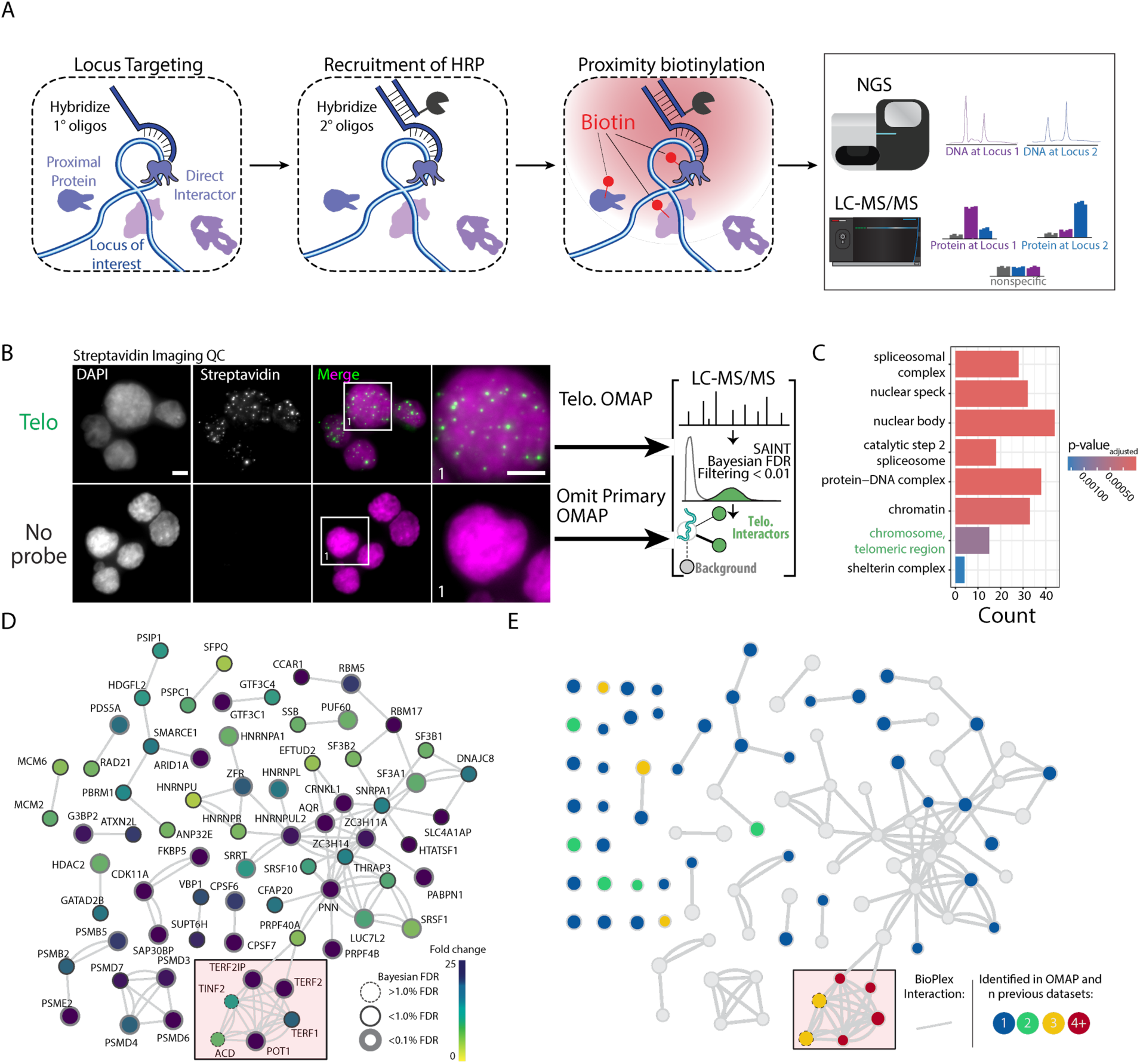
Overview of DNA O-MAP workflow and label-free quantitative proteomics analysis of telomeres. A) Schematic of DNA O-MAP. B) Fluorescent microscopy data showing the observed patterns of DNA (DAPI, left) and *in situ* biotinylation detected by staining with fluorescent streptavidin conjugates (middle, left) and overview of telomere targeted DNA O-MAP experiment. C) Significant gene sets identified by the Gene Set Enrichment Analysis of the proteins enriched by the telomere probe. D) DNA O-MAP telomeric proteins mapped onto the BioPlex interaction network^32,33^. The red box highlights shelterin complex proteins. Nodes are colored by the fold-enrichment compared to a no-primary-probe control shown in B, excluding unconnected nodes. E) Telomeric proteins observed in five previous datasets (PICh, C-BERST, CAPLOCUS, CAPTURE, BioID) superimposed onto Figure 1E, colored by the number of prior datasets where the protein was present and including unconnected nodes. Scale bars, 5 µm.

### DNA O-MAP reveals the organization of the telomeric proteome

To demonstrate that O-MAP can successfully purify proteins from genomic viewpoints, we selected human telomeres for initial testing (**Figure 1A**). Mammalian telomeres are several kilobases of tandemly repeated arrays of 5’-TTAGGG-3’ hexamers with terminal 3’ single-stranded overhangs at the ends of chromosomes^30^. Telomeric DNA is specifically bound by a proteinaceous cap that protects the natural chromosome ends from being recognized as damaged DNA—the shelterin complex^31,32^. Shelterin is a six-subunit complex, which is comprised of the telomeric repeat-binding factor 1 (TERF1), telomeric repeat-binding factor 2 (TERF2), protection of telomeres protein 1 (POT1), adrenocortical dysplasia protein homolog (ACD), TERF2-interacting protein 1 (TERF2IP), and TERF1-interacting nuclear factor 2 (TINF2). Due to the unique telomeric sequence and characteristic DNA structure, the shelterin proteins accumulate at the ends of the chromosomes. Accordingly, this well-defined set of proteins has been widely accepted as goalposts for a successful locus-proximal enrichment experiment^12,13,16^. In the near-diploid HCT-116 cells, telomeres have an average length of 5.6 kb and their cumulative length approximates 0.017% (∼500kb) of the human genome (**Table S4**)^33^. Compared to other repetitive elements in the human genome, telomeres are relatively short in HCT-116 cells and thus serve as a rigorous test case for DNA viewpoints of around 500 kb in aggregate across the genome.

We performed a DNA O-MAP experiment in which we either targeted telomeric DNA or omitted the primary hybridization probe (negative control). We purified biotinylated proteins from <60 million cells in three technical replicates followed by imaging of biotinylation and identification of proteins using label-free, MS1-based quantitative proteomics. By streptavidin staining, the punctate fluorescence pattern of biotin-labeled biomolecules closely mimicked telomere FISH, whereas we did not observe patterns of these puncta in the negative control samples (**Figure 1B**). From our label-free proteomics analysis, we identified 163 proteins as significantly enriched at telomeres. As expected, gene set enrichment analysis^34^ identified significant enrichment of telomeric chromosomal components, chromatin, and protein-DNA complexes (**Figure 1C-D**). Importantly, we identified all six shelterin proteins in the telomere sample and these proteins were completely absent from the control samples. Of the six shelterin proteins, four (TERF1, TERF2, TERF2IP, POT1) passed stringent false-discovery rate control while ACD and TINF2 did not due to low spectral intensity.

To benchmark DNA O-MAP, we compared the full set of telomeric proteins to proteins observed in five established telomeric datasets (PICh, C-BERST, CAPLOCUS, CAPTURE, BioID)^12,14,16,35,36^ (**Figure 1E**). DNA O-MAP captured both previously observed telomeric interacting proteins (shelterins) as well as telomere associated proteins (ribonucleoproteins). We identified multiple heterogeneous nuclear ribonucleoproteins (hnRNPs) previously annotated as telomere-associated, including HNRNPA1 and HNRNPU. HNRNPA1 has been demonstrated to displace replication protein A (RPA) and directly interact with single-stranded telomeric DNA to regulate telomerase activity^37–39^. HNRNPU belongs to the telomerase-associated proteome^40,41^ where it binds the telomeric G-quadruplex to prevent RPA from recognizing chromosome ends. We mapped DNA O-MAP enriched telomeric proteins to the BioPlex protein interactome^42,43^ and observed that in addition to capturing proteins from previously observed telomeric datasets (**Figure 1E**), DNA O-MAP enriched for protein-protein interactors with previously observed telomeric proteins. Previous data found RBM17 and SNRPA1 at telomeres, and in BioPlex these proteins interact with three SF3 proteins (SF3A1, SF3B1, SF3B2). Though they were not identified in previous telomeric proteome datasets, all three of these SF3 proteins were enriched in the DNA O-MAP telomeric data. Furthermore, through interactions with G-quadruplex binding factors, these SF3 proteins are regulators of telomere maintenance^44^.

### DNA O-MAP quantitatively compares nuclear- and mitochondrial-targeted DNA proximal proteomes

We next evaluated the utility of DNA O-MAP to quantitatively measure proteins associated with specific genomic loci. We integrated sample multiplexing quantitative^29,45–47^ proteomics downstream of DNA O-MAP to enable spectral quantification of all samples simultaneously (**Figure 2A**). In our experimental design, we measured the proteomes at three well-characterized DNA loci with distinct protein occupants in the human genome and a no-primary probe negative control: 1) telomeres, 2) peri-centromeric alpha satellite repeats, 3) the mitochondrial genome, 4) no primary probe negative control (**Figure 2B**). Centromeres are epigenetically defined chromosomal loci where kinetochore proteins assemble for spindle microtubule attachment to ensure equal chromosome segregation during cell division^48,49^. Human centromeres are located within the AT-rich alpha satellite repeats, which are higher-order repeats composed of 171-base-pair monomeric units^50,51^. Due to the sequence independence of centromeres, we utilized a previously described probe^28,52^ that targets a subset of alpha satellite repeats to represent centromeres, hereafter denoted as the ‘Pan Alpha’ probe (**Table S4**). The predicted genome-wide binding profile^53^ of the pan-alpha probe closely overlaps with centromeres and covers an estimated 35 Mb (**Figure S1**). Mitochondria are intracellular organelles of eukaryotic cells with their own genome (mtDNA). The mtDNA is a circular double-stranded DNA molecule of about 16.6 kb, located in the mitochondrial matrix associated with the inner membrane^54,55^. In HCT-116 cells, mtDNA copy number has been reported to range from 310–677^56^.

**Figure 2.**
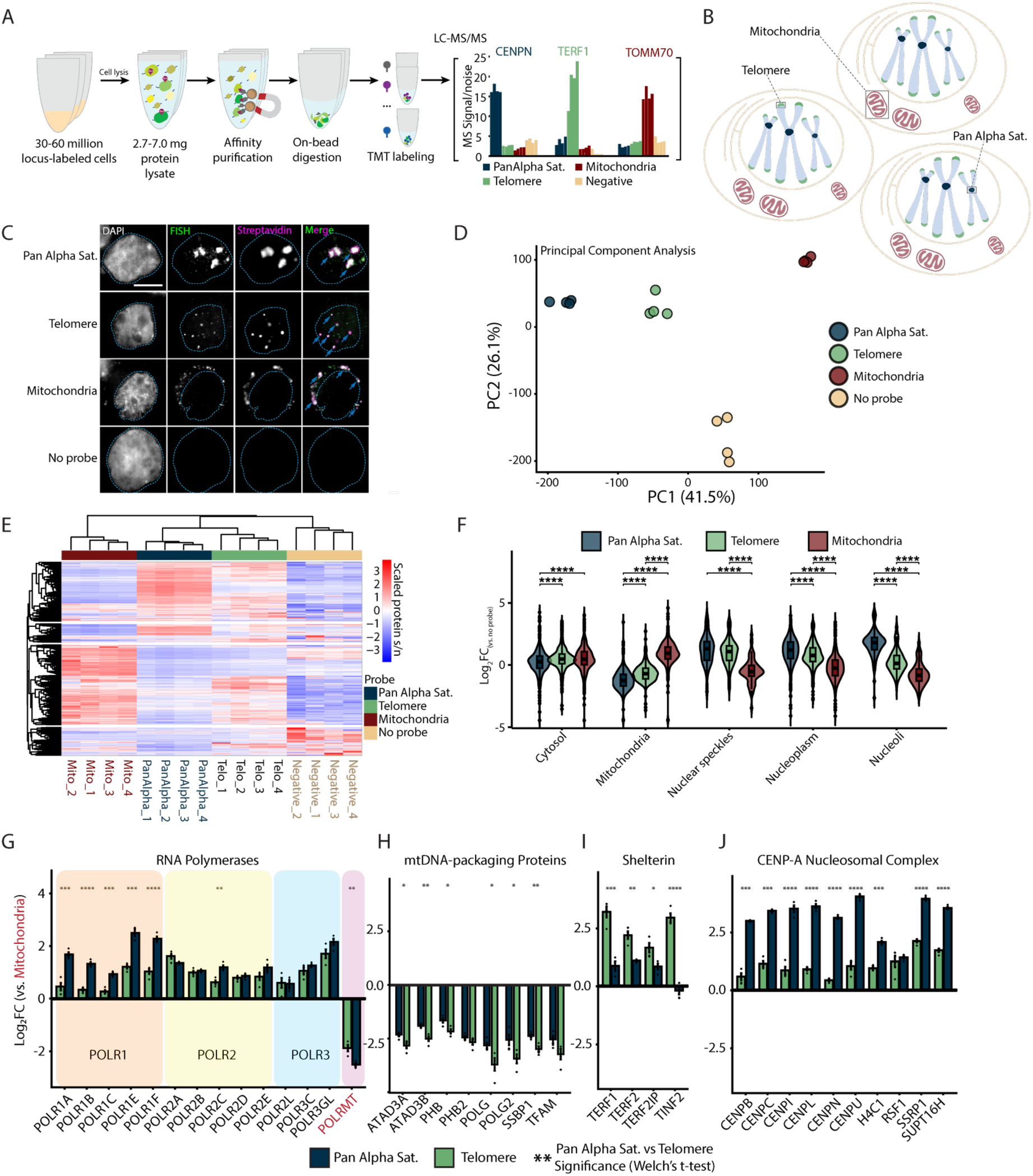
DNA O-MAP reveals distinct features of the sub-proteomes at peri-centromeric alpha satellites, telomeres, and the mitochondrial genome. A) Workflow of DNA O-MAP integrated with sample multiplexing quantitative proteomics. B) Schematic of the three DNA loci examined in the TMT16plex experiment: peri-centromeric alpha satellites, telomeres, and mitochondrial genomes. C) Co-localization of DNA FISH and the streptavidin staining of the proteins biotinylated by DNA O-MAP targeting the peri-centromeric alpha satellites, telomeres, and mitochondrial genomes. Scale bar: 5 µm. D) Principal component analysis of scaled intensities of proteins enriched by the pan-alpha probe, telomere probe, mitochondrial genome oligo pool, and no-primary-probe control. E) Unsupervised hierarchical clustering of scaled intensities of proteins enriched by the pan-alpha probe, telomere probe, mitochondrial genome oligo pool, and no-primary-probe control. F) Log2 fold change of proteins compared to no-primary-probe control, grouped by HPA subcellular location. Significance calculated based on Welch’s t-test for pairwise comparisons (****: p-value <0.0001). G–J) Log2 fold change of proteins compared to mitochondrial probe enriched proteins for the RNA Polymerases (G), mtDNA nucleoid packaging proteins^58^ (H), Shelterin (I), and CENP-A nucleosomal complexes (J). Significance calculated based on Welch’s t-test for pairwise comparisons (p-value: *<0.05, **<0.01, ***<0.001, ****<0.0001).

To determine the localization of biotinylation using the new oligos and oligo pools, we performed DNA O-MAP in human HCT-116 cells with a co-hybridization of both fluorescent oligos and HRP oligos in order to observe fluorescent *in situ* hybridization (FISH) and *in situ* biotinylation signals in the same cell. Biotinylation patterns of the pan-alpha, telomere, and mtDNA probes showed strong concordance with FISH (**Figure 2C**). To quantify the local proteomes corresponding to each of these biotinylated patterns, we prepared replicate (n=4) samples for each probe and control (**Figure S2**). Comparison of FISH and biotin signal revealed that DNA O-MAP labeling was highly specific to each of the telomeric, centromeric, and mitochondrial FISH signals (**Figure S3**). After *in situ* HRP-mediated labeling, we performed thermal reversal of fixation of cells prior to lysis, enrichment of biotinylated proteins^57^, tryptic digestion, and labeling with isobaric TMTpro barcodes^29^. We confirmed that artifactual lysine acylation due to cellular fixation with PFA did not widely affect TMTpro labeling of peptides as only 1.38% of lysine-containing peptides were acylated with fixative. Notably, the fixative was not present during protease digestion, and thus all peptide N-termini were available for TMTpro labeling and quantitation, irrespective of lysine modification status.

In total we quantified 3,055 proteins across all four targeted and control samples (**Figure 2D–E**). We observed consistent proteome enrichment for replicate analyses with O-MAP by principal component analysis and correlation analyses (**Figure 2D–E, S2**). Based on Human Protein Atlas annotations^59^, we observed significant enrichment of mitochondrial proteins with the mtDNA-probe proteomes and proteins from nuclear locations such as nuclear speckles, nucleoplasm, and nucleoli enriched by the telomere and pan-alpha probes (**Figure 2F, S4**). Notably, the pan-alpha probe enriched proteins from the nucleoli, consistent with the known nucleoli-centromere associations^60^ and chromosomal passenger complex member AURKB, consistent with the centromeric localization of AURKB in early mitosis to ensure faithful chromosome segregation^61,62^ and the localization of chromosomal passenger complex members to pericentromeric heterochromatin^63,64^. We also observed pericentromeric enrichment of spindle and chromosomal segregation associated proteins TPX2^65^ and KIF20A^66^ (**Figure S4, S5**).

Next, we explored the enrichment of several multi-unit protein complexes across the examined loci. To dissect the differences between enriched proteomes for each probe, we chose a subset of proteins of interest and measured the fold change of the two nuclear targets compared to mitochondria. RNA Polymerase I,II,III subunits were all higher in the nuclear probes than mitochondria, however in contrast to RNA Polymerase II and III, POLR1 proteins are significantly enriched in pan-alpha compared to telomere (**Figure 2G**). This enrichment is likely due to clustering of centromeres around nucleoli^67,68^, the location of ribosomal RNA synthesis by RNA Polymerase I. Conversely, mitochondrial RNA Polymerase POLRMT abundance was significantly lower in the nuclear probe proteomes compared to the mitochondrial probe proteome (log_2 Pan-Alpha Sat./Mito._= -2.51; log_2 Telomere/Mito._= -1.88). Similarly, we observed enrichment of mtDNA-packaging nucleoid components^58^ with the mtDNA probes (TFAM, SSBP1, POLG, POLRMT, Lon, ATAD3A/B, and PHB/PHB2; **Figure 2G–H**). As above, we observed consistent enrichment of shelterin components at telomeres (**Figure 2I**). We also observed CENP-A nucleosomal complexes enriched in the pan-alpha proteomes (**Figure 2J**). Histones were enriched with our nuclear probes and a subset (H2A1C, H2AX, and H4C1) were significantly enriched by the pan-alpha probe compared to the telomere probe (**Figure S5**). We also observed enrichment of catenins CTNNB1 and CTNND1 at telomeres (**Figure S4**). The transcription factor CTNNB1 has been observed at the transcriptional start site of h*TERT* where it regulates h*TERT* expression^69^. The h*TERT* gene is located in the subtelomeric region of chromosome 5 (chr5:1,253,167-1,295,068) and expressed in HCT-116 cells^70^. Collectively, these results highlight the subcompartment sensitivity of DNA O-MAP to distinguish differential compartment components even for ubiquitous chromatin constituents like histones.

### DNA O-MAP uncovers DNA-DNA interactions from non-repetitive DNA loci

Beyond repetitive regions in the human genome, we explored whether DNA O-MAP can recover material from single-copy DNA intervals. To this end, we designed an experiment in which we performed *in situ* biotinylation followed by chromatin extraction, affinity purification, and sequencing (**Figure 3A**). The human genome is folded into thousands of chromatin loops where two loci on the same chromosome are tethered to each other (**Figure 3B**). The anchors of the loops are bound by the insulator protein CTCF. The ring-shaped cohesin protein complex is thought to often stall at CTCF-bound sites while dynamically moving along the genome, creating contact domains of preferential DNA-DNA interaction^71^. These contacts between chromatin loop anchors have been captured genome-wide with *in situ* Hi-C^72^. Normally present in two copies per genome, these 20–25 kb loop anchor intervals are considerably less abundant than telomeres.

**Figure 3.**
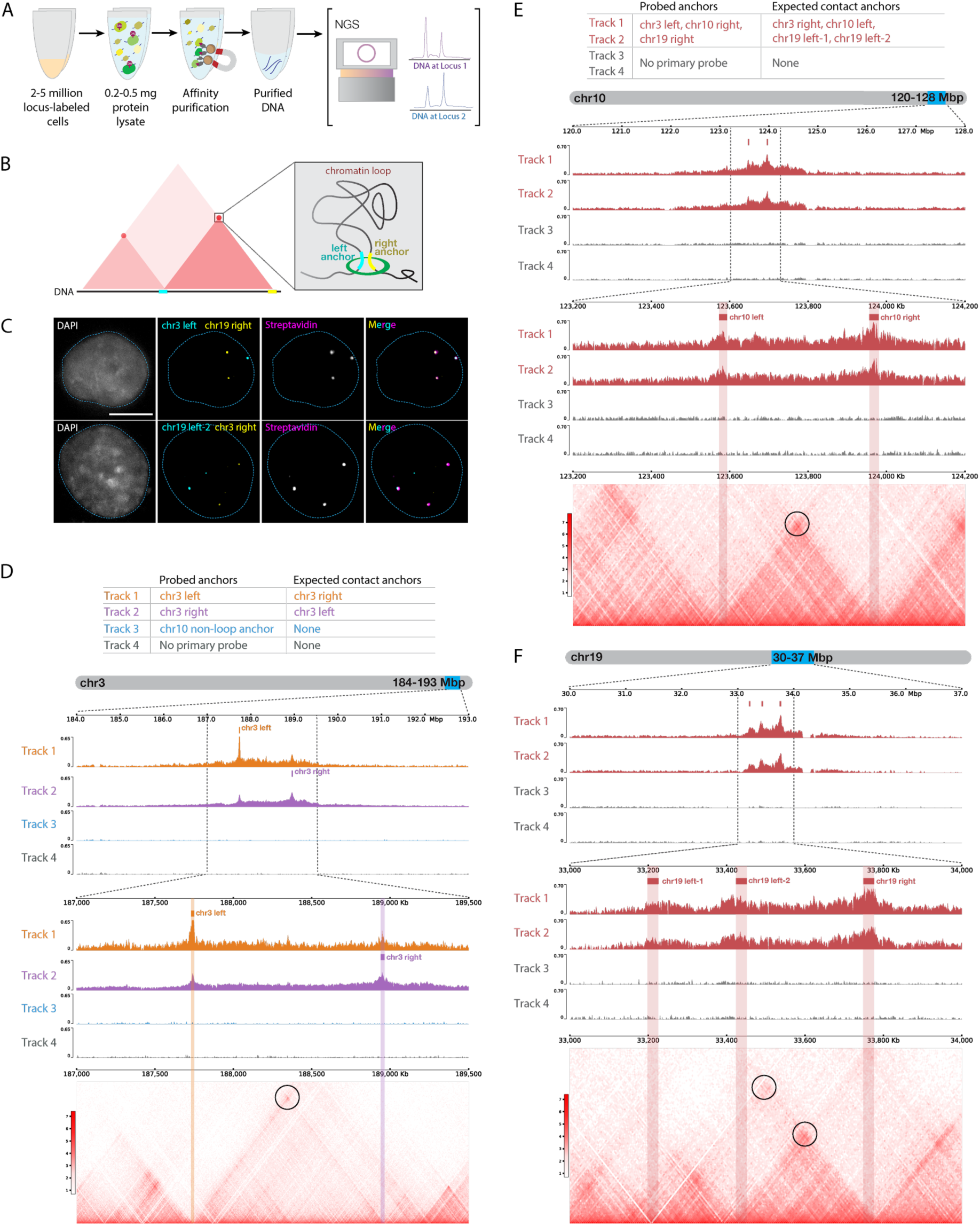
DNA O-MAP efficiently labels single-copy chromatin loop anchors. A) Workflow of DNA O-MAP integrated with biotin purification sequencing. B) Schematic of a pair of chromatin loop anchors on a hypothetical Hi-C map and 3-dimensional space. C) DNA FISH and the streptavidin staining of the proteins biotinylated by DNA O-MAP targeting anchors of chromatin loops on chromosome 3 and chromosome 19. D) Table listing the three anchors (Track 1-3) and no-primary-probe control (Track 4) biotinylated by DNA O-MAP and their expected anchors in contact in each track (top). Desthiobiotin purification sequencing signals across the 9-Mb region on chromosome 3 corresponding to the chr3 chromatin loop (middle). Desthiobiotin purification sequencing signals and pairwise contact map at 5-kb resolution across the 2.5-Mb region on chromosome 3 corresponding to the chr3 chromatin loop. Black circle on the contact map indicates the presence of a loop (bottom). E) Table listing the three chromatin loop anchors (Track 1-2) and no-primary-probe controls (Track 3-4) biotinylated by DNA O-MAP in duplicates and their expected anchors in contact in each track (top). Desthiobiotin purification sequencing signals across the 8-Mb region on chromosome 10 corresponding to the chr10 chromatin loop targeted (middle). Desthiobiotin purification sequencing signals and pairwise contact map at 5-kb resolution across the 1-Mb region on chromosome 10 corresponding to the chr10 chromatin loop. Black circle on the contact map indicates the presence of a loop (bottom). F) Desthiobiotin purification sequencing signals across the 7-Mb region on chromosome 19 corresponding to the chr19 chromatin loops targeted (top). Desthiobiotin purification sequencing signals and pairwise contact map at 5-kb resolution across the 1-Mb region on chromosome 19 corresponding to the chr19 chromatin loops. Black circles on the contact map indicate the presence of loops (bottom).

We first evaluated whether DNA O-MAP can specifically biotinylate loop anchors with microscopy by a co-hybridization of both fluorescent oligos and HRP oligos at four anchors: chr3 left (chr3:187,729,712-187,749,712), chr3 right (chr3:188,939,711-188,964,711), chr19 left-2 (chr19:33,425,000-33,450,000), and chr19 right (chr19:33,750,000-33,775,000). DNA O-MAP specifically biotinylated the biomolecules proximal to these small DNA intervals, as observed in the co-localizing patterns of FISH and streptavidin staining in the same cells (**Figure 3C**). We next evaluated whether DNA O-MAP could recover the DNA interactions originally discovered by Hi-C. We targeted a pair of intervals with high contact frequency—chr3 left and chr3 right anchors, one non-looping interval (chr10:123,187,984-123,207,984), and a no-primary-probe control. We performed DNA O-MAP to biotinylate these DNA intervals, subjected the labeled cells to chromatin solubilization and desthiobiotin purification, and sequenced the eluate DNA. As expected, all three probed DNA intervals were highly enriched compared with other genomic regions, indicating efficient purification of the loci (**Figures 3D, S6A**). Furthermore, chr3 left and chr3 right anchors reciprocally recovered each other, indicating that DNA O-MAP was able to recover known DNA interactions mediated by proteins. In contrast, the non-looping chr10 anchor did not enrich any other peak other than itself (**Figure S6B**). Lastly, in the cells that received no primary oligos, no pronounced enrichment was observed genome wide (**Figure S6B**).

To examine the multiplexability and reproducibility of DNA O-MAP, we simultaneously targeted three chromatin loop anchors: chr3 left, chr10 right (chr10:123,957,984-123,977,984), and chr19 right anchors in duplicates and subjected the cell pellets to purification and DNA sequencing. All three targeted anchors, chr3 left, chr10 right, and chr19 right anchors were successfully enriched (**Figures 3E–F, S7A**), whereas no pronounced enrichment was observed in the no-primary-probe controls genome-wide (**Figure S7B**). Furthermore, chr10 left (contacting chr10 right), chr19 left-1, and chr19 left-2 (both contacting chr19 right) were also efficiently recovered, accurately matching the Hi-C contact maps and the signals from two replicates was consistent (**Figure 3E–F, S8**). We confirmed this by distance-dependent normalization previously developed for proximity-labeling methods (**Figure S8**)^73^. These imaging and genomics data demonstrate that DNA O-MAP is capable of labeling small, single-copy DNA intervals with high specificity.

### DNA O-MAP uncovers differential protein enrichment from non-repetitive loci

Given our success with recovering DNA-DNA contact information from unique loci, we explored whether DNA O-MAP can recover the local proteome from single-copy DNA intervals as we did earlier with repetitive intervals targeting telomeres, alpha satellite repeats, and the mitochondrial genome. To investigate this, we utilized the *HOXA* and *HOXB* gene clusters as targets. The *HOX* subgroup of the homeobox family of genes has been well described for their roles in determining body plan formation, cell identity, and has been implicated in susceptibility to oncogenesis^74–78^. Here we designed probes to target 83kb and 81kb for *HOXA* and *HOXB*, respectively. We performed DNA O-MAP targeting either *HOXA*, *HOXB*, or no primary hybridization probes from 50 million human K562 cells across six replicates for *HOXA* and *HOXB* and five replicates for no primary (**Figure 4A**). We also performed DNA O-MAP in human K562 cells with co-hybridization of both fluorescent oligos and HRP oligos to demonstrate locus-targeted biotinylation and FISH signal within the same cell. Biotinylation patterns for both *HOXA* and *HOXB* showed strong concordance with the FISH signal (**Figure 4B**). Identification of biotinylated proteins was carried out using label-free quantitative proteomics. From our label-free proteomics analysis, we identified 42 proteins that were significantly differentially enriched between the *HOXA* and *HOXB* loci (**Figure 4C, S9A**). Of the differentially enriched proteins, both HDAC3 and TCF12 were scored as enriched in the *HOXB* labeled sample compared to *HOXA*. This agrees with ENCODE^79^ ChIP-seq data in K562 cells, with more called peaks for these proteins at *HOXB* than *HOXA* (**Figure 4D**). Additionally, SMARCB1 was scored as enriched in the *HOXA* labeled sample compared to *HOXB* and ENCODE^79^ ChIP-seq data showed enrichment of SMARCB1 at *HOXA* over *HOXB* (**Figure 4D**). SWI/SNF factors such as SMARCB1 have been described to interact with the *HOX* genes^80^, but differential enrichment has not been well documented.

**Figure 4.**
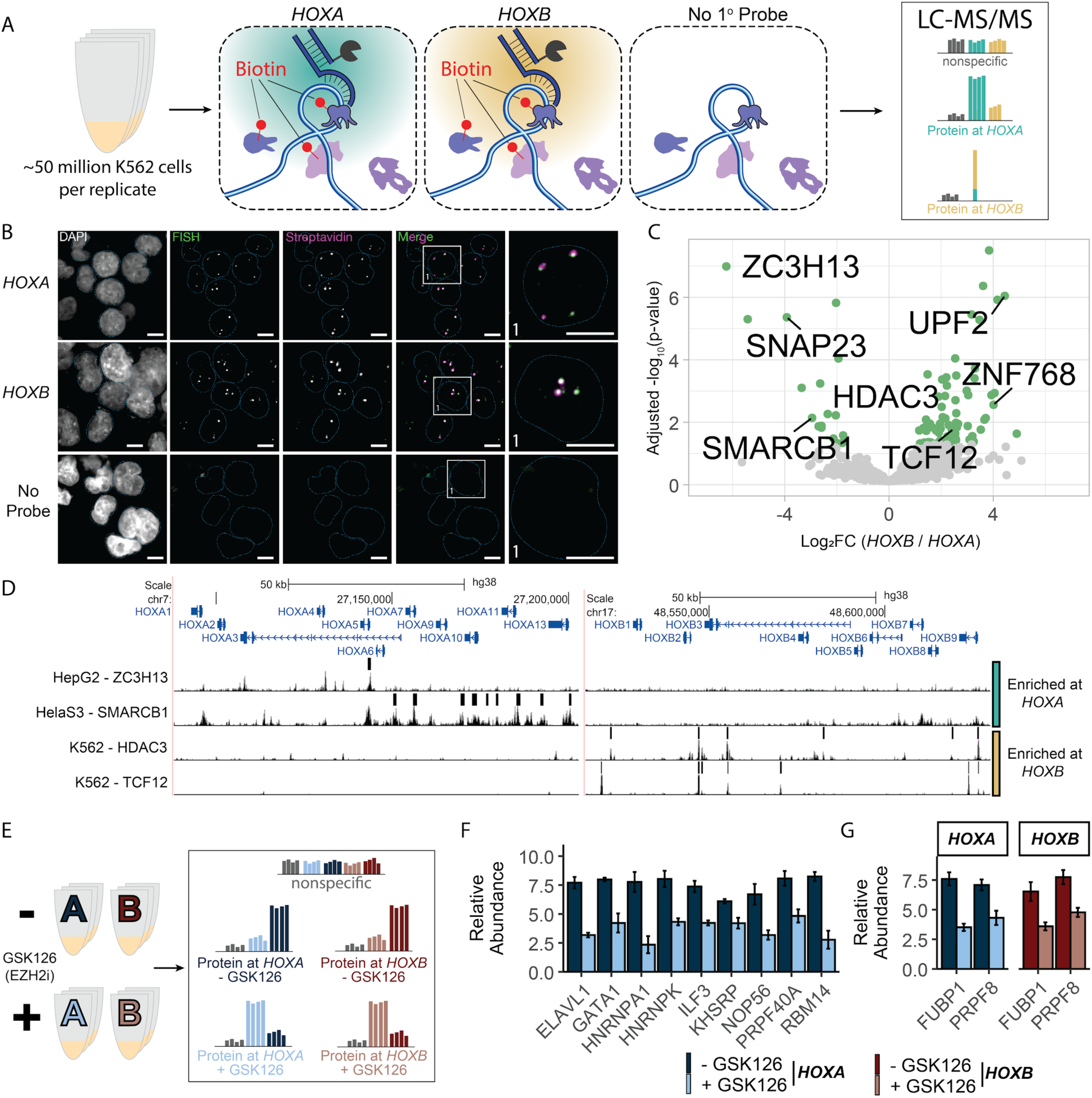
DNA O-MAP efficiently identifies the local proteome of the *HOXA* and *HOXB* gene clusters. A) Schematic of DNA O-MAP being applied to the *HOXA* and *HOXB* gene clusters for identification of differentially enriched proteins. B) Representative images depicting overlap of FISH and Streptavidin labeling at *HOXA* and *HOXB* loci. C) Volcano plot of proteins identified at *HOXA* and *HOXB* loci. Each dot represents a single protein with proteins of interest called out in black. Green dots indicate proteins that passed significant enrichment thresholds with an absolute Log2 Fold Change greater than 1 (2-fold change) and corrected p-value < 0.05. D) ENCODE ChIP-seq data showing peak calls and p-values at *HOXA* and *HOXB* loci for selected enriched proteins, ZC3H13, SMARCB1, HDAC3, and TCF12. E) Schematic depicting the use of GSK126 with DNA O-MAP. F) Bar chart showing proteins with significantly altered abundance following treatment with GSK126 at *HOXA*. G) Bar chart showing proteins with significantly altered abundance at both *HOXA* and *HOXB* following treatment with GSK126 (Welch’s t-test, corrected p-value < 0.05).

To further test the sensitivity and application of our non-repetitive DNA O-MAP approach at the *HOX* genes, we performed a chemical perturbation analysis targeting *HOXA* and *HOXB* in the presence or absence of enhancer of zeste homolog 2 (EZH2) inhibition with GSK126. In differentiated cells, such as K562s, the *HOX* genes are silenced by the Polycomb Repressive Complex 2 (PRC2). The main catalytic component of PRC2 is EZH2, which is responsible for trimethylating histone 3 at lysine 27 (H3K27me3)^81,82^. Deposition of H3K27me3 has been well characterized as a mark of transcriptional repression associated with PRC2 activity. Additionally, EZH2 inhibition has become a prominent target in cancer therapies^83,84^. For the perturbation analyses, we used sample multiplexing quantitative proteomics^29,45,46^ and DNA O-MAP to quantify the local proteomes in response to drug inhibition (**Figure 4E**). We prepared replicate (n=3) samples for each probe following treatment with either the EZH2 inhibitor GSK126 or DMSO control. In total, we identified 11 proteins at *HOXA* and 8 proteins from *HOXB* with significantly altered enrichment following EZH2 inhibition (**Figure 4F, G, S9B**), many of which are RNA-interacting proteins, such as ELAV1, RMB14, ILF3, HNRNPK, HNRNPA1, PRPF40A, and PRPF8. These data are consistent with recent evidence that suggests that long non-coding RNA-interacting proteins are critical for the establishment of PRC2 repressed loci^85,86^. Several proteins we identified also have established connections to PRC2 repression of the *HOX* genes, such as GATA1 which has been suggested to mediate the switch to non-canonical PRC2 function^87^. These data demonstrate that DNA O-MAP has sufficient sensitivity to detect proteomes at non-repetitive genomic loci and measure inhibitor-induced changes to the proximal proteome at sub-megabase, non-repetitive genomic regions (∼80 kb).

### Establishing an on-plate hybridization-biotinylation workflow for DNA O-MAP

During the development of DNA O-MAP, we refined an on-plate workflow on cells adhered to glass bottom plates (**Figure 5A**). We began with adherent cells grown in 6-well plates, using 3 wells per replicate per condition with each well yielding 0.75 - 1 million cells. The cells are subsequently fixed (4% PFA) in order to be compatible with DNA ISH. Multiple plates can be processed in parallel to enable multiple replicates per experiment. We note that the on-plate version of the protocol further reduces the total number of cells required per replicate by ∼10-fold compared to the in-solution protocol.

**Figure 5.**
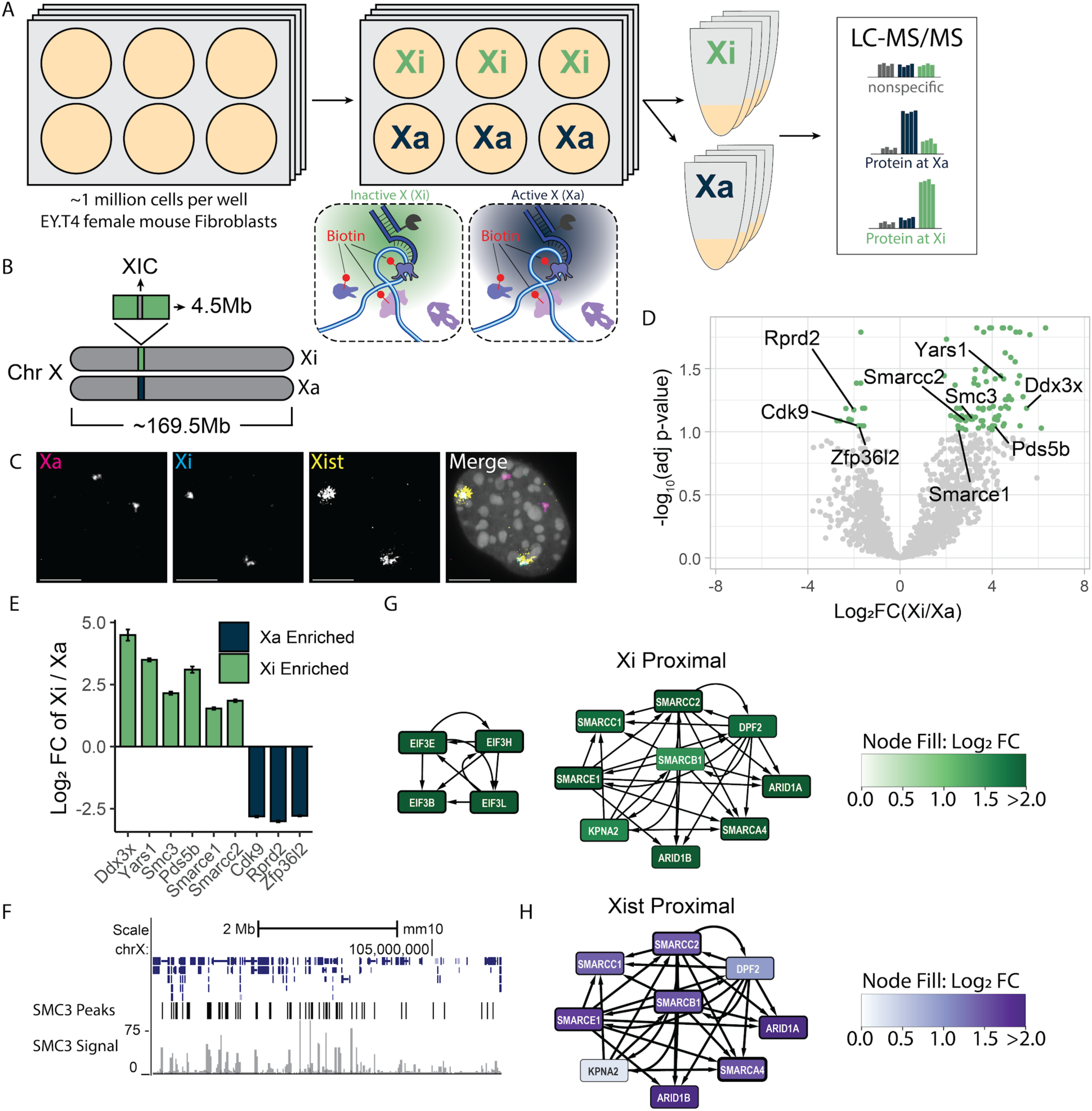
DNA O-MAP elucidates the homolog-resolved chromosome X proteome. A) Schematic of DNA O-MAP being applied to Xi and Xa for identification of differentially enriched proteins. B) Schematic showing the region of the X chromosome targeted by our primary hybridization probes. C) Representative images depicting overlap of Xist FISH and Xi Streptavidin labeling while spatially differentiated from Xa FISH. Scale bars are 10 uM. D) Volcano plot of proteins identified at Xa and Xi. Each dot represents a single protein with proteins of interest called out in black text. Green dots indicate proteins that passed significant enrichment thresholds with an absolute Log2 Fold Change greater than 1 (2-fold change) and corrected p-value < 0.1. E) Bar chart showing example proteins with significant enrichment at Xi in green and at Xa in blue. Corrected p-value < 0.1. F) ENCODE ChIP-seq data in mouse fibroblast cells at our targeted region of chromosome X for SMC3. G) Protein interaction networks of EIF and SWI/SNF complexes enriched at Xi. Node width is a function of corrected p-value and node color is a function of enrichment (Log2 Fold Change). H) Protein interaction network of the SWI/SNF complex from previously published RNA O-MAP of Xist. Node width is a function of -Log10(p-value) and node color is a function of enrichment (Log2 Fold Change).

### DNA O-MAP detection of homolog specific proteomes

To further understand the level of sensitivity we can achieve when investigating the local proteome of non-repetitive loci, we explored whether DNA O-MAP can recover material in a homolog specific manner. To this end, we designed primary hybridization probes that differentially target either the active or inactive X-Chromosome homolog (Xa and Xi, respectively). The Xa-Xi paradigm is a particularly interesting variable homolog target as the Xi is largely transcriptionally silenced through cooperative activity of the non-coding RNA Xist and H3K27me3 by the PRC, along with other factors, in direct opposition of the Xa^85,86,88–90^. Thus, these two homologs present a vastly different chromatin environment. To individually target Xi and Xa, we used differential single nucleotide polymorphisms (SNPs) between the maternal and paternal homologs in EY.T4 female mouse fibroblast cells that were identified by homolog resolved genome sequencing^91,92^. In doing so we were able to design two sets of primary hybridization probes to target the same 4.5Mb region on either Xi or Xa straddling the X inactivation center (**Figure 5B**). The approach of leveraging SNPs for homolog specific probe binding has been previously validated for use with other ISH-based methods^92,93^, but has not been previously applied to proximity labeling or proteomics approaches For validation of our differential labeling approach we performed FISH for Xist and Xa alongside streptavidin labeling of Xi. Labeling patterns of Xi and Xist overlapped as expected while remaining spatially distinct from Xa FISH (**Figure 5C**). We performed a DNA O-MAP experiment in which we targeted either Xa or Xi across four replicates. Each replicate consisted of three wells of a 6-well plate each containing 0.75 - 1 million cells (**Figure 5A**). Of note, in contrast to the previous experiments, here we did not release our adherent cells and all steps were carried out with cells fixed to glass-bottom 6-well plates prior to lysis. From our label-free proteomics analysis, we identified 96 proteins that were significantly differentially enriched at either Xi or Xa (**Figure 5D, S10**). The Xa showed enrichment of proteins associated with a more active transcriptional pattern in comparison to Xi, such as RPRD2, ZFP36l2, and CDK9 (**Figures 5D & E**). For the Xi in comparison to Xa, we observed enrichment of protein interactors and complexes^42,43^, including the EIF3 family, members of the cohesin complex Smc3 and Pds5b, and SWI/SNF factors. Both the cohesin and SWI/SNF complexes have been previously shown to interact with the inactive X chromosome^94^ and enrichment of SWI/SNF factors was consistent with recent work measuring the proteome of Xist, the Xi-resident long non-coding RNA^27^ (**Figure 5G**). Interestingly, we also see enrichment of Ddx3x, an X-linked gene previously implicated as a mediator of sex-based differences in neurodevelopment and disease^95^. Here we show that DNA O-MAP was able to obtain homolog-specific, differentially enriched local proteomes from a 4.5Mb, non-repetitive, region on the X chromosome from less than 3 million cells per replicate.

## Discussion

By combining the versatility of hybridization-based genome targeting with the robustness of proximity biotinylation, DNA O-MAP offers a scalable approach to study DNA proximal proteomes of a specific locus. The hybridization-biotinylation workflow allows for efficient processing of samples and is compatible with both proteomic and genomic readouts. Integration with multiplexed quantitative proteomics enables simultaneous analysis of multiple loci or conditions, increasing data completeness and throughput. Label-free analysis of the telomeres shows strong concordance of labeling with in-situ hybridization and recapitulates previous similar proteomic datasets. Our tri-locus experiment was able to differentiate proteins with a quantitative profile suggesting general nuclear location from those specifically associated with telomeres and peri-centromeres. DNA O-MAP can target single-copy loci, as evidenced by the chromatin loop anchor, *HOXA* and *HOXB* gene cluster, and X-chromosome targeting experiments, which enables the study of a wide range of DNA proximal proteomes. The ability to detect differences in these proteomes is exemplified by the application of DNA O-MAP to the *HOXA* and *HOXB* loci following EZH2 inhibition and to the X-chromosome in a homolog specific manner.

O-MAP has now been shown to be a highly flexible technology for the exploration of biomolecular interactions with RNAs^27^ and DNA loci. Using oligos to target the DNA locus, DNA O-MAP can be theoretically adapted for use in any sample type amenable to *in situ* hybridization, including cultured cells, tissue sections, and primary tissue samples^28,53,96^. As the purification tag is decoupled from the probe oligos, labeled chromatin fragments can undergo stringent washes to achieve efficient purification with minimal background. Moreover, without the need to genetically modify the biological system at hand, the probes in this dataset alone could be used to explore telomeric remodeling in cancer cells^36^, spindle-associated proteome dynamics at the pericentromere^97^, and molecular drivers of hetero-or euchromatin formation^98^ at nearly any locus in the human genome (O-MAP probes can feasibly cover >99% of the human genome)^53,96^. Additionally, we show that this method is able to detect DNA-DNA contacts through detection of biotinylated loop anchors. Our approach functions similarly to 4C^99^, however, the biotin labeling of contacts does not rely on pairwise ligation events. Thus, detection of contacts through DNA O-MAP will vary in the sampling of DNA-DNA contacts in comparison.

DNA O-MAP has several current limitations. First, fixation can modify proteins and must be accounted for when labeling is performed in fixed cells. Second, DNA O-MAP captures proximal proteins for a given genomic locus. This means measured proteins will be both direct interactors and also proteins that are reproducibly within close physical proximity to the locus. This distinction has been described previously^27,100^, but is generally important for interpretation of DNA O-MAP proteomics results and design of future DNA O-MAP experiments. This is due, at least in part, to the long-lived nature of the biotin-phenoxy radical species generated by peroxidase-based proximity labeling methods. Third, DNA O-MAP may miss proteins at specific loci if the protein is especially low abundance, highly post-translationally modified, or otherwise challenging to detect by mass-spectrometry-based proteomics. Finally, DNA O-MAP reports more proteins than those that are ‘specific’ to a given locus. For example, all nuclear targeting probes identified nuclear, but non-specific, proteins like histones. An important aspect of the quantitative approaches used here is that proteins are measured as differentially abundant at a given loci in comparison to other loci or across treatments.

By taking a comparative quantitative approach, we remove the need to pre-define the local context of probe localization, but experimental design is critical and novel interactors may require further validation to confirm their co-localization at a given locus (e.g., with imaging/FISH). With developments in automation and instrument sensitivity, DNA O-MAP has the potential to expand to post-translational modifications and be used for large-scale chromatin perturbation screens. We anticipate that DNA-OMAP will have broad utility for research questions seeking to understand the intricate relationships between DNA sequence, chromatin structure, and cellular function.

## Methods

### Cell culture and fixation

Colorectal cancer HCT-116 cells were grown in ATCC-formulated McCoy’s 5A Medium Modified (ATCC 30-2007). EY.T4 mouse fibroblasts were grown in ATCC-formulated DMEM (Gibco 11965-092). Human lymphoblast K562 cells were grown in ATCC-formulated RPMI (Gibco 11875-093). All cell lines were supplemented with 10% fetal bovine serum and 100 U/ml Penicillin-Streptomycin and grown at 37°C in a humidified atmosphere of 5% CO_2_. For each HCT 116 purification, 20 million cells were seeded into one T-500 flask (Thermo Scientific 132867) to culture for 36-48 hours to reach 90–120 million cells. Before collection, cells were briefly rinsed once with Dulbecco’s phosphate buffered saline (DPBS) and then incubated with 25 ml of TrypLE Express Enzyme (Gibco 12604-021) at 37°C for two minutes or until loosely attached. For each K562 purification, 25 million cells were seeded into one T-500 flask to culture for 48 hours to reach 100 million cells. Before collection, cells were briefly rinsed once with DPBS. For inhibition of EZH2, K562 cells were grown and harvested, as described above, in medium supplemented with either 5uM GSK126 (Tocris 6790) or DMSO for 48 hours and then collected. For both HCT 116 and K562 purifications, the cell suspension was collected into two 50 ml conical tubes and the T-500 flask was rinsed with DPBS. The wash was combined with the cell suspension and centrifuged at 300 G for 5 minutes. After a DPBS wash to remove remaining TrypLE, cells were fixed in 4% paraformaldehyde (wt/vol) (Electron Microscopy Sciences 15710) in PBS in suspension at room temperature for 10 minutes with rotation, followed by 125 mM Glycine quenching for 5 minutes at room temperature with rotation and 15 minutes on ice. Fixed cells were collected by centrifugation at 350G for 5 minutes, and stored in fresh DPBS at 4°C until liquid-phase hybridization. Fixed cells were used within 3-5 days.

For each EY.T4 purification, 300,000 cells were seeded into each well of a glass-bottom 6-well plate (Cellvis P06-1.5H-N) to culture for 24 hours to reach 750,000 - 1 million cells per well. Cells were briefly rinsed once with DPBS then fixed in 4% paraformaldehyde (wt/vol) (Electron Microscopy Sciences 15710) in PBS in at room temperature for 10 minutes. Fixed cells were then washed with DPBS three times for 5 minutes each. Fixed cells were stored in fresh DPBS at 4*C until solid-phase hybridization. Fixed cells were used within 5 days.

### Primary oligo probes

Primary oligos targeting the human alpha satellite repeat and telomere were purchased as individually column-synthesized DNA oligos from Integrated DNA Technologies. Probe sets targeting *HOXA* (chr7:27092311 - 27175959), *HOXB* (chr17:48527413 - 48608584), mtDNA (chrM:1-16,569), chr3 left anchor (chr3:187,729,712-187,749,712), chr3 right anchor (chr3:188,939,711-188,964,711), chr10 non-looping anchor (chr10:123,187,984-123,207,984), chr10 right anchor (chr10:123,957,984-123,977,984), and chr19 right anchor (chr19:33,750,000-33,775,000) were designed using PaintSHOP^96^ and ordered in oPool format from Integrated DNA Technologies. Homolog specific chromosome X (chrX:100,254,241 - 102,428,950, 102,601,850 - 104777629) targeting probes were designed using an in-house computational pipeline and ordered in oPool format from Integrated DNA Technologies. More than 300 primary oligos were designed to cover each single-copy DNA interval to ensure a sufficient number of probes at the locus for FISH. The sequences of the oligo and oligo pools used are listed in Supplementary Dataset 1.

### Oligo library synthesis

Amplification of the *HOXA*, *HOXB*, Xi, and Xa targeting primary oligo libraries was performed as follows. 20 ng µl−1 of oligo library in 10 mM Tris, pH 8.0 was amplified by PCR. The reaction mix contained 34 µl of dH2O, 10 µl 5× Phusion HF Buffer, 1.5 µl of 10 mM dNTP Mix, 1.5 µl of 10 µM F primer, 1.5 µl of 10 µM R primer, 1.0 µl of resuspended oligo pool and 0.5 µl of Phusion DNA Polymerase (2 U µl−1). The thermal cycler program was as follows; 95 °C for 3 min, followed by 12 cycles of 98 °C for 20 s, 60 °C for 15 s, 72 °C for 15 s then 72 °C for 1 min followed by a 4 °C hold. This PCR product was purified using a Zymo DNA Clean and Concentrator-5 (DCC-5) kit according to the manufacturer’s protocol. 20 pg µl−1 of the first PCR product was prepared as a template for the second PCR. The second PCR mix contained 27 µl of dH2O, 10 µl of 5× Phusion HF Buffer, 1.5 µl of 10 mM dNTP Mix, 5.0 µl of 10 µM F Primer, 5.0 µl of 10 µM R Primer, 1.0 µl of diluted DNA template and 0.5 µl of Phusion DNA Polymerase (2 U µl−1). The thermal cycler program was repeated, but for 18 cycles instead of 12. The second PCR product was purified as before. RNA was synthesized using the NEB HiScribe T7 Quick High Yield RNA Synthesis Kit with a modified reaction mix containing 8 µl of dH2O, 2.5 µl of diluted DNA template, 15.0 µl of NTP Buffer Mix, 3 µl of T7 RNA Polymerase Mix and 1.5 µl of RNaseOUT. The reaction was carried out for 16 hours at 37 °C followed by a 12*C hold. The DNA template was then digested using DNase I (M0303) by adding 2uL (4 units) of DNase I and 50uL of Ultra Pure Water (UPW) (10977-015) to each reaction. The reverse transcription reaction contained 45 µl of synthesized RNA, 15 µl of 5× RT Buffer, 10.5 µl of 10 mM dNTP Mix, 1 µl of 100 µM RT Primer, 1.5 µl of RNaseOUT and 2 µl of Maxima H Minus Reverse Transcriptase (200 U µl−1) and 20ul of UPW for a total volume of 75 µl. The reactions were incubated at 50 °C for 2 h, 85 °C for 5 min, then a 4*C hold. RNA templates were degraded by adding 37.5 ul of 0.5M EDTA and 37.5 ul of 1M NaOH to each reaction and incubated at 95*C of 10min followed by a 4*C hold. Final ssDNA probe was purified using a Zymo DNA Clean and Concentrator-100 (DCC-100) kit according to the manufacturer’s protocol for oligo purification.

### Primer exchange reaction (PER)

To extend primary oligos with PER concatemers, reactions were set up as previously described^101^ in 100 ul-volume containing 10 mM MgSO4, 300 uM dATP/dCTP/dTTP mix, 100 nM Clean.G hairpin, 80 U/ml Bst DNA Polymerase, Large Fragment (NEB M0275L), 1 uM hairpin, and 1 uM primary oligos in PBS. To verify the length of primary oligos, the reactions were assessed with denaturing polyacrylamide gel electrophoresis. Primary oligos extended to 300-500 nucleotides were used in hybridizations downstream. Unpurified reactions were dehydrated using vacuum concentrators and stored dry at -20°C until hybridization.

### In-solution hybridization and biotinylation of cell pellets

HCT 116 and K562 oligo hybridizations were performed on cells in solution for the cost-effectiveness of primary and secondary oligos. Fixed cells were split into 6e7 cell aliquots in 1.5 ml microcentrifuge tubes. All washes and buffer exchanges were performed as follows: centrifuging at 350G for 3.5 minutes or until pelleted, pouring away used buffers from the pellets, adding new buffers, and gentle shaking or low speed vortexing to dislodge cell pellets into tiny clusters or cell suspensions for incubations or washes. Cells in fresh wash buffer were rotated on a low speed nutator for 5 minutes.

Cells were rinsed once with fresh phosphate buffer saline (PBS), and permeabilized in PBS-0.5% TritonX-100 (Sigma T8787) for 10 minutes with nutation. After a PBS-0.1% Tween20 (PBS-T) (Sigma T2287) wash, permeabilized cells were incubated in 0.1 N hydrochloric acid (HCl) for 5 minutes. After a PBS-T wash to remove acid, cells were incubated in PBS-T-0.5% hydrogen peroxide to block endogenous peroxidases. After a 2X saline sodium citrate-0.1% Tween20 (2X SSC-T) wash to remove acid, cells were incubated in 2X SSC-T-50% formamide for 20 minutes at 60°C on a Thermomixer C dry block (Eppendorf 2231001005). Cells were exchanged into primary hybridization buffer (Hyb1) comprising 2X SSC-T, 50% (vol/vol) formamide, 10% (wt/vol) dextran sulfate, 0.4 μg/ul RNAse A, and ∼1 μM extended primary oligos (resuspended dry, unpurified PER reactions). The cell-Hyb1 mixture was distributed into PCR strip tubes at 1e7-1.5e7 cells in 100 μL volumes. The cells were denatured and primary oligos were hybridized to the genome in the PCR strip tubes in a thermocycler using the cycling protocol: 78°C 3 minutes, 37°C `∞` incubating overnight for more than 18 hours. The next day, cells were rinsed with 60°C 2X SSC-T into 1.5 ml microcentrifuge tubes, followed by two 2X SSC-T buffer exchanges to remove residual Hyb1. Cell pellets were then washed in 1 ml 2X SSC-T at 60°C, followed by two two-minute washes in 2X SSC-T at room temperature. Fully washed cell pellets were exchanged into 1 ml PBS, and then exchanged into 100 nM secondary HRP oligo that map to the PER concatemer sequence on the primary oligo (custom synthesis by Integrated DNA Technologies or Bio-Synthesis Inc) in PBS. Secondary hybridization was performed at 37°C with nutation for one hour. Cell pellets underwent three 5-minute washes in 1 ml PBS-T at 37°C with nutation. Fully washed cells were incubated in 5 uM desthiobiotin tyramide (Iris Biotech LS-1660) and 1 mM hydrogen peroxide in PBS-T for 5 minutes at room temperature with nutation. To quench the HRP activity, biotinylated cells were washed twice in 10 mM sodium ascorbate and 10 mM sodium azide in PBS-T for 5 minutes at room temperature with nutation. Quenched cells were washed with PBS to remove residual sodium azide. After sampling cells for quality control, the cell pellets were stored dry in -80°C until chromatin solubilization and affinity purification.

### On-plate hybridization and biotinylation

EY.T4 oligo hybridizations were performed on cells fixed to glass bottom 6-well plates, with all washes being performed in 1mL. Cells were rinsed once with fresh phosphate buffered saline (PBS), and permeabilized in PBS-0.5% TritonX-100 (Sigma T8787) for 10 minutes. After a PBS-0.5% Tween20 (PBS-T) (Sigma T2287) wash, permeabilized cells were incubated in 0.1 N hydrochloric acid (HCl) for 5 minutes. After a PBS-T wash to remove acid, cells were incubated in PBS-T-0.5% hydrogen peroxide to block endogenous peroxidases. After a 2X saline sodium citrate-0.1% Tween20 (2X SSC-T) wash to remove acid, cells were incubated in 2X SSC-T-50% formamide for 20 minutes on a heat block in a 60°C water bath. Cells were exchanged into 500uL per well of primary hybridization buffer (Hyb1) comprising 2X SSC-T, 50% (vol/vol) formamide, 10% (wt/vol) dextran sulfate, 0.4 μg/ul RNAse A, and ∼250 nM extended primary oligos. The cells were denatured on a heat block in a water bath at 78°C for 3 minutes. The cells were then hybridized with primary oligo at 37°C overnight with nutation for about 24 hours. The next day, cells were rinsed with 60°C 2X SSC-T four times for five minutes each, followed by two two minute 2X SSC-T buffer exchanges at room temperature to remove residual Hyb1. Fully washed cells were then washed with 1 ml PBS followed by 100 nM secondary HRP oligo that map to the concatemer sequence on the primary oligo in PBS. Secondary hybridization was performed at 37°C with nutation for one hour. Cells underwent three 5-minute washes in PBS-T at 37°C. Fully washed cells were incubated in 5 uM desthiobiotin tyramide (Iris Biotech LS-1660) and 1 mM hydrogen peroxide in PBS-T for 5 minutes at room temperature. To quench the HRP activity, biotinylated cells were washed twice in 10 mM sodium ascorbate and 10 mM sodium azide in PBS-T for 5 minutes at room temperature. Quenched cells were washed with PBS to remove residual sodium azide. The cells were stored dry at 4°C until chromatin solubilization and affinity purification.

### Microscopy-based quality control assays for hybridization and biotinylation

We routinely sample cells along the workflow of preparing AP-MS or NGS samples to monitor the specificity of primary oligo hybridization. To assess the quality of primary oligo hybridization for the liquid-phase prepared cells, we sampled roughly 5% of fully washed cells from primary hybridization to a new 1.5 ml tube. Cells were incubated with 400 nM fluorescent oligos in PBS at 37°C for an hour with nutation. Hybridized cells underwent three washes in 1 ml PBS-T at 37°C with nutation to remove unbound fluorescent oligos. Washed cells were immobilized on glass slides with Slowfade Gold Antifade Mountant with DAPI (Thermo Fisher S36938) and coverslips for confocal imaging of FISH signal.

We assessed the quality of biotinylation specificity for all samples entering the proteomics or genomics workflow. For liquid-phase, roughly 5% of fully quenched cells were sampled into a new 1.5 ml tube and incubated with 0.5-1 μg/ml Alexa Fluor 647-streptavidin (Thermo Fisher S32357) in PBS-T, 1% bovine serum albumin at 37°C for 30 minutes with nutation. Stained cells underwent four washes in 1 ml PBS-T at 37°C with nutation to remove unbound Alexa Fluor 647-streptavidin conjugate. Washed cells were immobilized on glass slides with Slowfade Gold Antifade Mountant with DAPI and coverslips for confocal imaging of Alexa-Fluor 647-streptavidin signals.

For the solid-phase workflow experiments in EY.T4 cells, we assessed primary oligo hybridization and biotinylation specificity simultaneously. A separate 6-well plate was prepared alongside the others that was reserved for quality control with one well per condition. After biotinylation and subsequent washes this plate was incubated with 0.5-1 μg/ml Alexa Fluor 647-streptavidin (Thermo Fisher S32357) in PBS-T, 1% bovine serum albumin at 37°C for 30 minutes with nutation. Stained cells underwent four washes in 1 ml PBS-T at 37°C with nutation to remove unbound Alexa Fluor 647-streptavidin conjugate. These cells were then incubated with 400 nM fluorescent oligos in PBS at 37°C for an hour. Hybridized cells underwent three washes in 1 ml PBS-T at 37°C with nutation to remove unbound fluorescent oligos. Washed cells were immobilized on glass slides with Slowfade Gold Antifade Mountant with DAPI and coverslips for confocal imaging of Alexa-Fluor 647-streptavidin and FISH signal.

### Confocal microscopy

Confocal imaging was performed using a Yokogawa CSU-W1 SoRa spinning disc confocal device attached to a Nikon ECLIPSE Ti2 microscope. Excitation light was emitted at 30% of maximal intensity from 405 nm, 488 nm, 561 nm, or 640 nm lasers housed inside of a Nikon LU-NF laser unit. Laser excitation was delivered via a single-mode optical fiber into the CSU-W1 SoRa unit. Excitation light was directed through a microlens array disk and a SoRa spinning disk containing 50 um pinholes to the rear aperture of a 100x N.A. 1.49 Apo TIRF oil immersion objective lens by a prism in the base of Ti2. Emission light was collected by the same objective and directed by a prism in the base of Ti2 back into the SoRA unit, where it was relayed by a 1x lens (conventional imaging) or 2.8x lens (super-resolution imaging) through the pinhole disk and then directed to the emission path by a quad-band dichroic mirror (Semrock Di01-T405/488/568/647-13X15X0.5). Emission light was then spectrally filtered by one of four single-bandpass filters (DAPI:Chroma ET455/50M; ATTO488: Chroma ET525/36M; ATTO565:Chroma ET605/50M; Alexa Fluor 647: Chroma ET705/72M) and focused by a 1x relay lens onto an Andor Sona 4.2B-11 camera with a physical pixel size of 11 um, resulting in an effective resolution of 110 nm (conventional), or 39.3 nm (super-resolution). The Sona was operated in 16-bit mode with rolling shutter readout and exposure times of 70-300 ms.

### FISH-biotinylation co-localization experiment

Fixed cells were split into 5e6 cell aliquots in 1.5 ml microcentrifuge tubes. Primary hybridization and washes were performed similarly to described in the in-solution hybridization and biotinylation of cell pellets with fewer cells. Fully washed cell pellets were exchanged into a secondary co-hybridization buffer containing 30 nM of fluorescent oligos and 100 nM of HRP-oligos in PBS, instead of solely HRP-oligos, for simultaneous hybridization of both species. After washes and biotinylation, the pellets were stained with 0.5-1 μg/ml Alexa-Fluor 647-streptavidin. Cells were immobilized on glass slides with Slowfade Gold Antifade Mountant with DAPI and coverslips for confocal imaging of both FISH and Alexa-Fluor 647-streptavidin signals.

### FISH-biotinylation co-localization quantification

To quantify the colocalization of signal in the FISH and Streptavidin channels, binary masks were generated for each image using a Gaussian blur (scipy.ndimage.gaussian_filter) and power-law transformation. The preprocessed image was then binarized using Otsu’s method (skimage.filters.threshold_otsu). Parameter optimization for the sigma value of the Gaussian blur and the exponent values of the power-law transformation were done via grid search. Each parameter set was used to generate a binary mask and scored based on the percentage of pixels labeled as foreground in the mask that exceeded the 90th percentile of pixel intensity values in the input image. Parameters were optimized for each image and channel independently. These binary masks were then used to calculate the labeling efficiency and specificity of the Streptavidin signal relative to the ground truth FISH signal and the Jaccard index of both masks. Labeling efficiency is defined as the percentage of pixels labeled as foreground in the FISH mask that were also labeled as foreground in the Streptavidin mask. Specificity is defined as the percentage of pixels labeled as foreground in the Streptavidin mask that were also labeled as foreground in the FISH mask. The Jaccard index was calculated by dividing the number of pixels in the intersection of the two masks by the number of pixels in their union.

### Affinity Purification and sample preparation for proteomics

For liquid-phase prepared samples, biotinylated cell pellets were removed from -80°C to thaw at room temperature. Each cell pellet was resuspended in roughly 0.9 ml of lysis buffer consisting of 1% SDS and 200 mM EPPS with protease inhibitors (Roche 11836170001). For the EY.T4 solid phase samples, 0.5 ml of lysis buffer was added directly to each well and cells were scrapped and collected in 1.5 ml tubes. The cell mixture was boiled at 95°C for 30 minutes. The boiled cell mixture was sonicated at 4°C using a Covaris LE-220 focused ultrasonicator with the following protocol: 300W peak incident power, 50% duty factor, 200 cycles per burst, with a treatment time of 420 seconds in 1-ml milliTUBEs with AFA fiber (Covaris 520135). The sonicated cell mixture was boiled for a second time at 95°C for 30 minutes. The boiled lysates were cleared by centrifuging at 21130 G for 30 minutes in an Eppendorf 5424 Microcentrifuge at room temperature. The supernatants were transferred to a fresh 1.5-ml tube. To prevent any remnants of cell debris, the supernatants were cleared for a second time by centrifuging at 21130 G for 30 minutes and the supernatants were transferred to a fresh 1.5-ml tube. The supernatants were stored in -80°C until protein quantification.

The cleared cell lysates were quantified using the Pierce BCA Protein Assay Kit (Thermo Fisher 23225). Pierce Streptavidin Magnetic Beads (Thermo Fisher 88817) were washed using 1% SDS, 200 mM EPPS lysis buffer three times before use. From each labeled cell pellet, 2.17 milligrams of protein was used to couple with 500 μg of streptavidin beads in a Protein Lo-Bind tube (Eppendorf EP022431081). The lysates were incubated with the bead slurry for one hour at room temperature with nutation allowing biotinylated proteins to bind. The coupled beads were collected and separated from the flow-through using a magnetic rack (Sergi Lab Supplies 1005a). After the flow-through was removed, the beads underwent the following washes: 2% SDS with 20 mM EPPS twice, 0.1 M Na2CO3, 2 M urea, and 1 M KCl with 20 mM EPPS twice. All washes were performed as follows: after immobilizing the beads on a magnetic rack for 5 minutes, the supernatant was removed, and the beads were resuspended in the new wash buffer and incubated for 5 minutes with nutation. Finally, the beads were rinsed once with 20 mM EPPS to remove the excess salt.

The washed streptavidin beads were resuspended in 50 μl of 5 mM TCEP, 200 mM EPPS, pH 8.5 for a 20-minute on-bead protein reduction. The proteins were alkylated on-bead using 10 mM iodoacetamide for one hour in the dark. Then DTT was added to the final concentration of 5 mM to quench the alkylation for 15 minutes. The beads were rinsed twice with 200 mM EPPS for on-bead digest. For liquid phase samples, assuming 20 μg of eluate protein, 200 ng LysC (Wako) was added to the beads in a 50-ul volume and incubated for 16 hours with vortexing. The next day, 200 ng of trypsin (Promega V5113) was added to the beads and incubated for six hours at 37°C at 200 rpm. For solid phase samples 20 ng of LysC and Trypsin was added and the samples were processed in the same way. After digestion, the peptide-containing supernatant was collected in a fresh 0.5-ml Protein Lo-Bind tube. The beads were rinsed once with 100 μl 50% acetonitrile, 5% formic acid and the wash was combined with the peptides. Peptides were desalted via the stop and go extraction (StageTip)^102^ method and dried in a vacuum concentrator. For label-free analysis of telomere-enriched samples, one sample consisted of HCT-116-Rad21-mAID cells^103^ . For samples intended to be multiplexed, dried, desalted peptides were reconstituted in 4 μl of 200 mM EPPS, pH 8.5. The peptides were labeled using 25 μg of TMTpro 16plex Label Reagents (Thermo Fisher A44520) at 33.3% acetonitrile for one hour at room temperature. The labeling reaction was quenched with the addition of 1 μl of 5% hydroxylamine and incubated at room temperature for 15 minutes. The pooled sample was acidified using formic acid and peptides were desalted using a StageTip cartridge. Peptides were eluted in 70% acetonitrile, 1% formic acid and dried by vacuum centrifugation

### Mass Spectrometry Data Acquisition Methods and Analysis

Samples were resuspended in 5% acetonitrile/2% formic acid prior to being loaded onto an in-house pulled C18 (Thermo Accucore, 2.6 Å, 150 μm) 30 cm column. Peptides were eluted over 180 min gradients running from 96% Buffer A (5% acetonitrile, 0.125% formic acid) and 4% buffer B (95% acetonitrile, 0.125% formic acid) to 30% buffer B. Sample eluate was electrosprayed (2700 V) into a Thermo Scientific Orbitrap Eclipse mass spectrometer for analysis. High field asymmetric waveform ion mobility spectrometry (FAIMS) was set at “standard” resolution, 4.6 L/min gas flow, and 3 CVs: −40/–60/–80 were used. Briefly, generally for both Label-free and TMTpro analyses, MS1 scans were collected at 120,000 resolving power with a 50 ms max injection time, and the AGC target set to 100%. Peaks from the MS1 scans were filtered by intensity (minimum intensity >5 × 103), charge state (2 ≤ z ≤ 6), and detection of a monoisotopic mass (monoisotopic precursor selection for peptides, MIPS). Dynamic exclusion was used with a duration of 90 s, repeat count of 1, mass tolerance of 10 ppm, and the “exclude isotopes” option checked. For each MS1, 8 data-dependent MS/MS scans were collected. MS/MS scans were conducted in the linear ion trap with the “rapid” scan rate, 50 ms max injection time, AGC target set to 200%, CID collision energy of 35% with 10 ms activation time (TMTpro) or HCD at 30% collision energy (Label-free), and 0.5 m/z (TMTpro) or 0.7 m/z (Label-free) isolation window. For TMTPro labelled samples, an MS3 scan was also included in the method. Unless otherwise noted in the methods, the real-time search filter was enabled^45^. Using a human fasta downloaded from Uniprot, fixed modifications for the TMTpro mass (+304.207146) were added to n-terminal residues and lysines. Carbamidomethly (+57.021464) was added for cysteines. Oxidation (+15.9949) was added as a variable modification on methionines. Missed cleavages were set to maximum of 1. “TMT mode” was enabled and thresholds of 1 and 0.05 for Xcorr and dCn respectively were used as minimums to trigger SPS-MS3 scans. SPS ions were set to 10 and MS3 scans were performed at a resolving power of 50,000, with an HCD collision energy of 45%, AGC of 200%, with a maximum injection time of 200 ms.

Label-free mass spectrometry data for MS1-based quantitation was analyzed with MSFragger^104^ search algorithm searched against a full human protein database for HCT 116 and K562 samples with forward and reverse protein sequences. EY.T4 samples were searched against a full mouse protein database with forward and reverse protein sequences. Fixed modifications included Carbamidomethyl (+57.021464) on cysteines. Variable modifications included oxidation (+15.9949) on methionine and formylation (+27.994915) on lysines. Peptides up to 2 missed cleavages were included. Peptide spectral matches and proteins were filtered to a 1% false discovery rate using Percolator^105^.

Multiplexed raw mass spectrometry data was analyzed using the Comet^106^ search algorithm, searched against a full human protein database with forward and reverse protein sequences (Uniprot 10/2020). Precursor monoisotopic peaks were estimated using the Monocle package. Fixed modifications included TMTpro (+304.207146) on n-terminal residues and lysines and Carbamidomethyl (+57.021464) on cysteines. Variable modifications included methionine oxidation (+15.9949) and lysine formylation (+27.994915). Peptides with up to 2 missed cleavages were included. Peptide spectral matches and proteins were filtered to a 1% false discovery rate using the rules of parsimony and protein picking. Protein quantification was done using signal-to-noise estimates of reporter ions. Samples were column normalized for total protein concentration. After filtering for contaminants, we performed a two-sided t-test comparing each O-MAP condition using Benjamini-Hochberg adjusted p values (i.e. q-values). Log_2_ fold changes of the mean of the biological replicates were also calculated for each biological condition. For the GSK treatment experiment, prior to graphing, the data were run through a custom R script using ComBat to correct for batch effects encountered when processing the samples. Human Protein Atlas^59^ subcellular locations were downloaded and the “main location” was assigned to each protein with a supported or enhanced reliability level. SAINT scores and interaction false discovery rates were calculated with the SAINTexpress software^107,108^. Significant hits were those with a SAINT calculated FDR less than 1%^109^. BioPlex interaction networks were accessed through the online BioPlex Explorer^110^. Networks were imaged using Cytoscape 3.10.02^111^. Protein complex members were accessed through CORUM^112^. Gene set enrichment analysis was performed with clusterProfiler^113^ and fgsea^114^ packages.

### Preparation of soluble chromatin for affinity purification followed by next generation sequencing

For confirmation of single-copy O-MAP labeling, loop anchor-biotinylated pellets of 10-20 million cells were removed from -80°C to thaw at room temperature. Each cell pellet was resuspended in an SDS lysis buffer consisting of 1% SDS and 200 mM EPPS with protease inhibitors. The cell mixture was sonicated at 4°C using a Covaris LE-220 focused ultrasonicator with the following protocol: 300W peak incident power, 15% duty factor, 200 cycles per burst, with a treatment time of 20-30 minutes in 130-μl microTUBEs with AFA fiber (Covaris 520077). After the samples had returned to room temperature, the sheared fixed chromatin was transferred to fresh 1.5-ml Protein Lo-Bind tubes and centrifuged at 21130 G for 10 minutes to pellet cellular debris. The supernatants were transferred to a new set of tubes. The cleared chromatin samples were quantified using the Pierce BCA Protein Assay Kit (Thermo Fisher 23225). Next, 50 μl of sheared chromatin was sampled for reverse crosslinking, DNA extraction, and gel electrophoresis to verify that a significant amount of DNA had been sheared to <700 base pairs. A sample of 10 μg sheared chromatin was reserved and stored at -20°C as immunoprecipitation input. 200 μg of chromatin was used to couple with 200 μg of streptavidin beads for one hour in a Protein Lo-Bind tube at room temperature with nutation. The coupled beads were collected and separated from the flow-through using a magnetic rack. After the flow-through was removed, the beads underwent the following washes:

○ 2% SDS with 20 mM EPPS
○ 2% SDS with 20 mM EPPS
○ High Salt Buffer containing 500 mM NaCl, 1 mM EDTA, 50 mM of HEPES pH7.5, 0.1% sodium deoxycholate, and 1% TritonX-100
○ LiCl Buffer containing 250 mM LiCl, 1 mM EDTA, 10 mM Tris-HCl pH 8.0, and 0.5% of IGEPAL CA-630
○ TE Buffer with 10 mM Tris and 1 mM EDTA
○ TE Buffer with 10 mM Tris and 1 mM EDTA

The washes were performed as follows: briefly spin and immobilize the beads on a magnetic rack, pipette out the supernatant as much as possible, resuspend the beads in 0.8 ml of wash buffer, and incubate for 5 minutes with nutation. The washed beads were resuspended in 300 ul of reverse crosslinking buffer containing 300 mM NaCl, 300 mM Tris-HCl pH 8.0, and 1 mM EDTA. Both the eluate beads and the input chromatin were incubated at 65°C for 16 hours for reverse crosslinking. The next day, 4 ul of 20 mg/ml proteinase K (Roche 3115836001) was added to the eluates and inputs and incubated at 50°C for 2 hours to cleave away proteins. The DNA was isolated from the mixture using phenol chloroform extraction followed by ethanol precipitation. Before sequencing library generation, the precipitated DNA was further purified using SPRI beads. The purified DNA was used to generate next-generation sequencing libraries using the NEBNext Ultra II DNA Library Prep Kit for Illumina (NEB E7645S) and NEBNext Multiplex Oligos for Illumina Index Primers Set 1 and 3 (NEB E7335S, E7710S) and PCR-amplified for 15 cycles. The sequencing libraries were quantified using the Qubit 4 fluorometer and library sizes were quantified using the D1000 ScreenTape assay (Agilent 5067-5582) on the TapeStation 4200 automated electrophoresis platform.

### DNA sequencing and data analysis

The libraries were mixed and sequenced pair-ended at 50-bp read length on an Illumina NextSeq 2000 sequencer to depths of 14.1-351.8 million reads per eluate sample and 3.14-16.45 millions reads per input sample using the NextSeq 1000/2000 P2 Reagents (100 Cycles) kit (Illumina 20046811). Reads were demultiplexed and adapters were removed using Cutadapt^115^. Trimmed reads were mapped to the reference genome (GRCh38) using Bowtie2 version 2.5.3 with the parameter -X 1000 keeping reads with a MAPQ>=30^116^. Duplicate reads were removed using Picard 3.1.1^117^. Eluate reads were normalized to input reads using DeepTools^118^ bamCompare with the following parameters: –binSize 20 –normalizeUsing BPM –smoothLength 60 – extendReads 150. Normalized data were visualized using Coolbox 0.3.9^119^.

To assess the targeting specificity of DNA O-MAP, O-MAP ChIP sequencing reads were aligned to a reference genome using Bowtie2. Duplicate reads were removed with Picard and Samtools. Enrichment scores were calculated using a bin-based method adapted from the TSA-seq protocol. The genome is divided into 100kb windows, and the enrichment score for each window is determined by the following formula: Enrichment Score = (Input reads in bin/Sum of input reads)/(Pull-down reads in bin/Sum of pull-down reads). To normalize the data and prevent division by zero in bins with no reads, a pseudocount equal to 10% of the total reads was added to each bin. Finally, the calculated enrichment scores were plotted against the log-scaled distance from the target regions.

## Data Availability

The mass spectrometry proteomics data have been deposited to the ProteomeXchange^120^ Consortium via the MassIVE with the data set identifier PXD054080. Sequencing data will be deposited to Gene Expression Omnibus before formal acceptance for publication. All primary data associated with the manuscript will be made available upon request.

## Author Contributions

Y.L., C.D.M., C.P.H., B.J.B., and D.K.S. conceived and designed the project. Y.L., C.D.M, C.P.H., M.K., T.A.P., R.F., C.H., S.A., A.F.T., and E.K. performed experiments. Y.L., C.D.M., C.P.H., N.J.L., and C.K.C. performed computational analyses. Y.L., C.D.M., C.P.H., B.J.B., and D.K.S. wrote the manuscript. All authors edited and reviewed the manuscript. D.M.S., B.J.B., and D.K.S. supervised the work.

## Competing Interest Statement

D.K.S. is a collaborator with Thermo Fisher Scientific, Genentech, Calico Labs, and AI Proteins. C.K.C., A.F.T., E.K., D.M.S., and B.J.B. have filed a patent application covering aspects of this work. B.J.B. is listed as an inventor on patent applications related to the SABER technology related to this work.

## Acknowledgements

We would like to thank members of the Shechner, Beliveau, and Schweppe labs for constructive feedback and technical assistance in assembling this work. We would also like to thank Drs. Jay Shendure, Shao-En Ong, Christine Quietsch, Emily Hatch, Gavin Ha, Celeste Berg, Christine Disteche, Andrew Stergachis, and Stanley Fields for helpful discussions of this work. We would like to acknowledge the following sources of support: R35GM137916 (BJB), R35GM150919 (DKS), the W.M. Keck Foundation (BJB, DKS), an Andy Hill CARE Distinguished Researcher Award (DKS), a Damon Runyon Dale Frey Award (BJB), a Cancer Consortium New Investigator Award (DKS), the Pew Charitable Trusts (DKS), 1R01GM138799-01 and 1R01HL160825-01 (DMS), T32GM007750 (to AFT and EEK), AHA 902616 (to EEK). Research reported in this publication was supported by the NHLBI under award number T32HL007093 (to CPH). This work was also supported by a Research and Education Training Fund Award (to CH) from the Center for the Multiplex Assessment of Phenotype at UW.

## Supplemental Information

**Figure S1.**
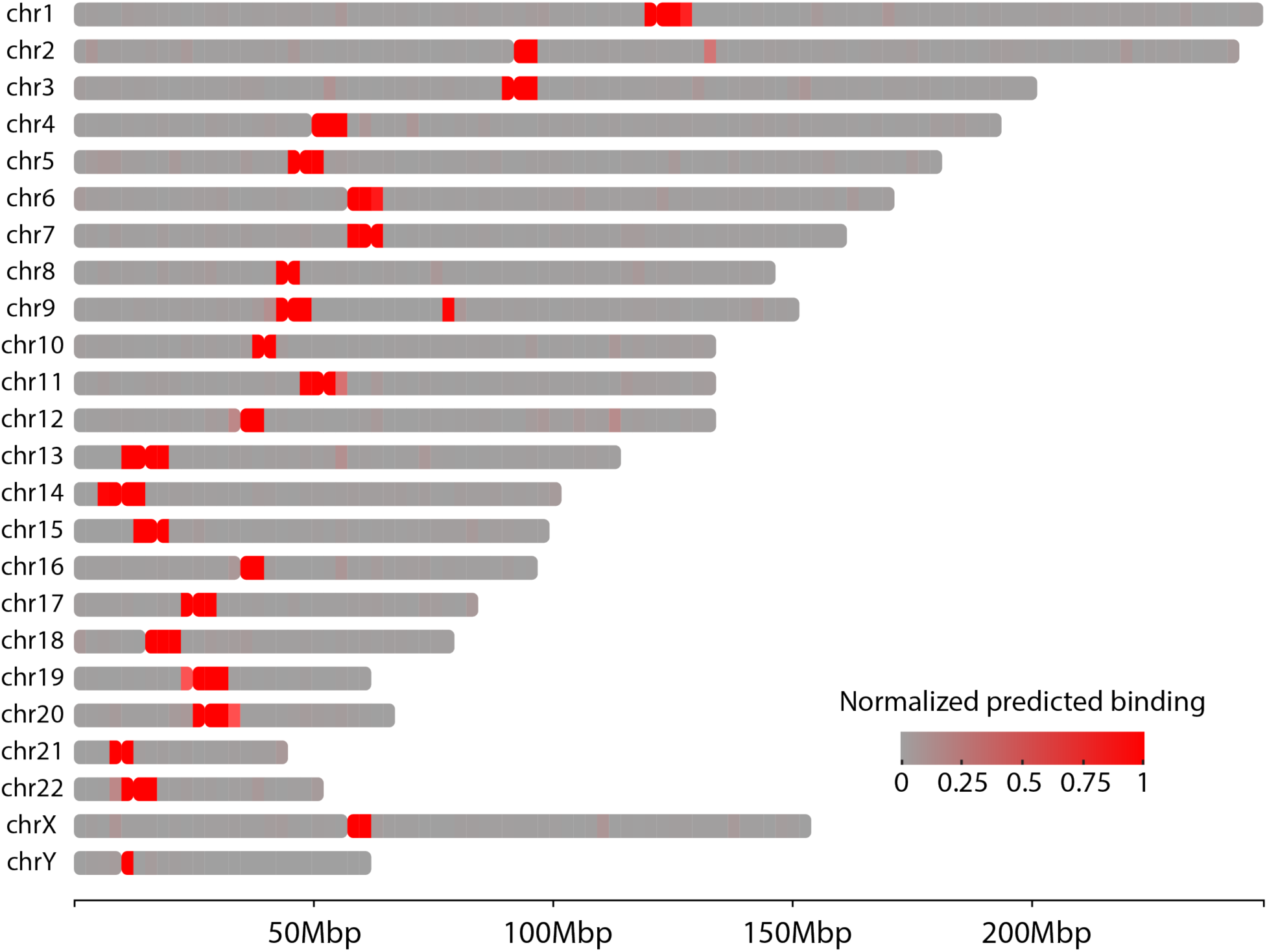
Predicted genome-wide binding profile of the pan-alpha probe. The intensity of red indicates the amount of predicted probe binding.

**Figure S2.**
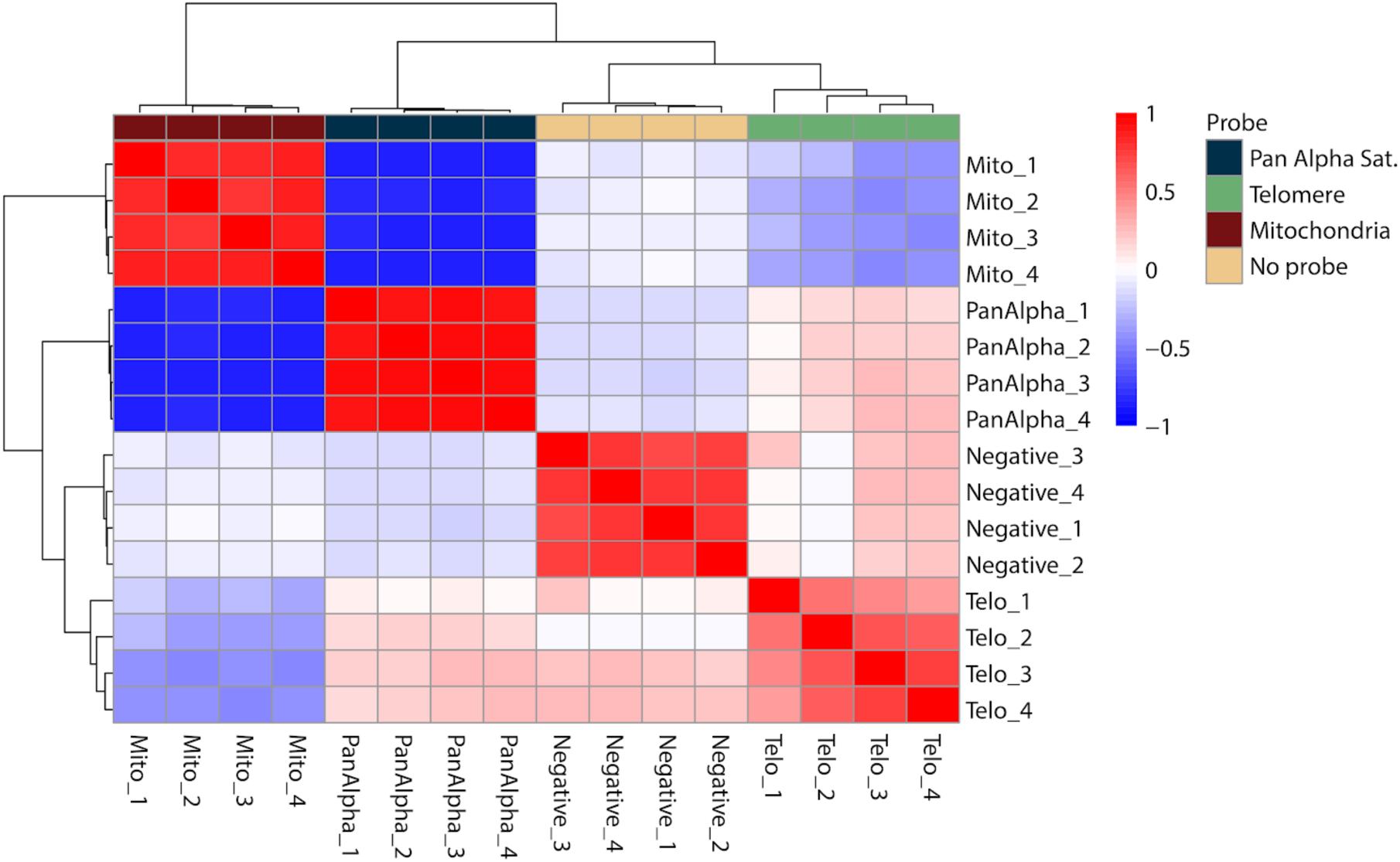
Replicate analysis of multi-target DNA O-MAP proteomics experiment. A) Pearson correlation coefficient of the raw protein intensity values for each replicate of the analysis with hierarchical clustering on the rows and columns.

**Figure S3.**
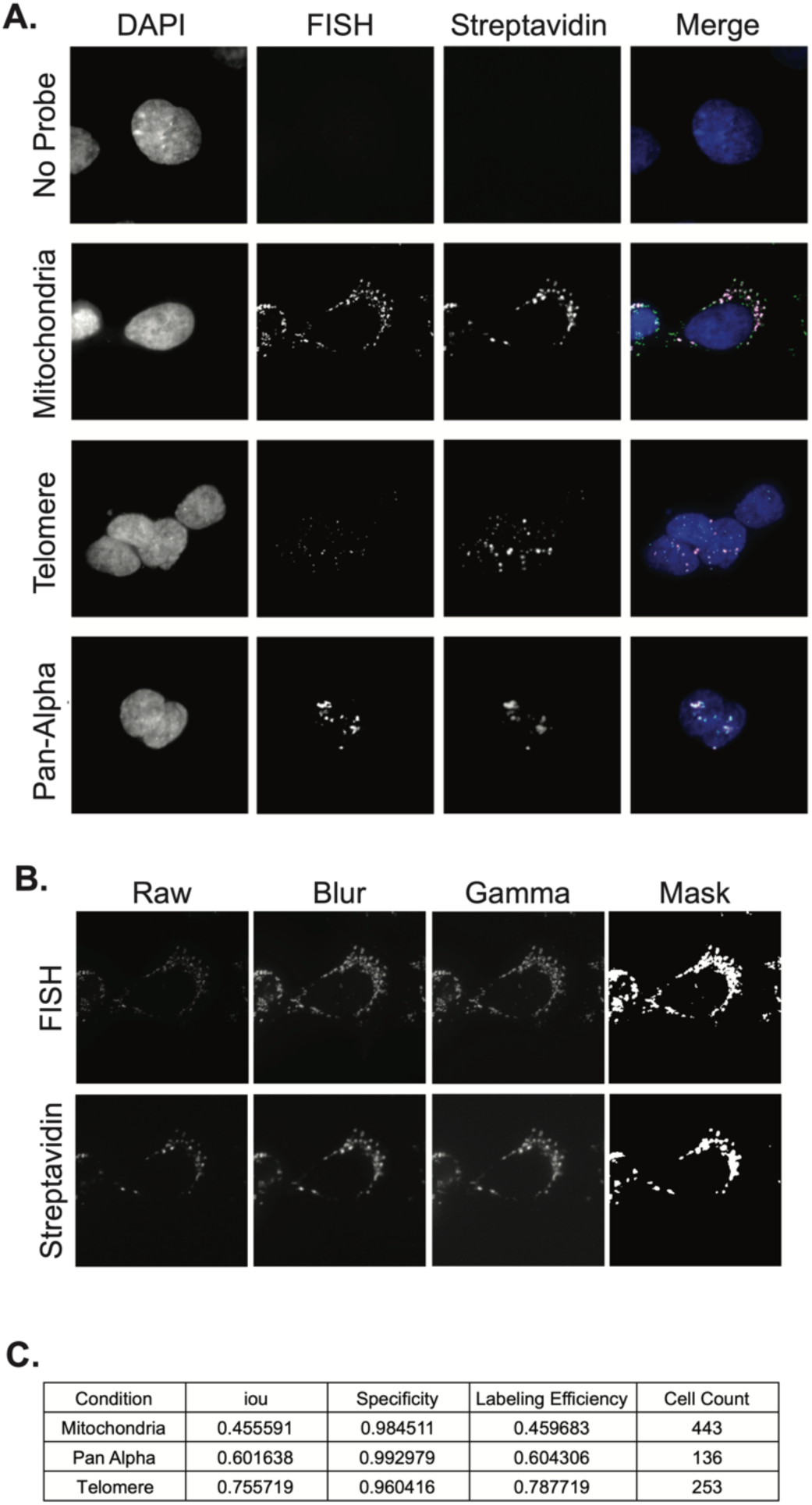
Quantification of DNA O-MAP labeling specificity and efficiency for pan-alpha, telomere, and mitochondria probes. A) Fluorescent microscopy data showing co-localization of DNA FISH and streptavidin staining of the proteins biotinylated by DNA O-MAP using no primary probe or targeting the mitochondrial genome, telomeres, or alpha satellites. B) Representative images for both DNA FISH and streptavidin stain from each stage of quantifying specificity and labeling efficiency for probes targeting the mitochondrial genome. C) Table of results for the mitochondrial genome, telomeres, and alpha satellites showing iou, specificity, labelling efficiency, and cell count of images quantified for DNA FISH and streptavidin staining.

**Figure S4.**
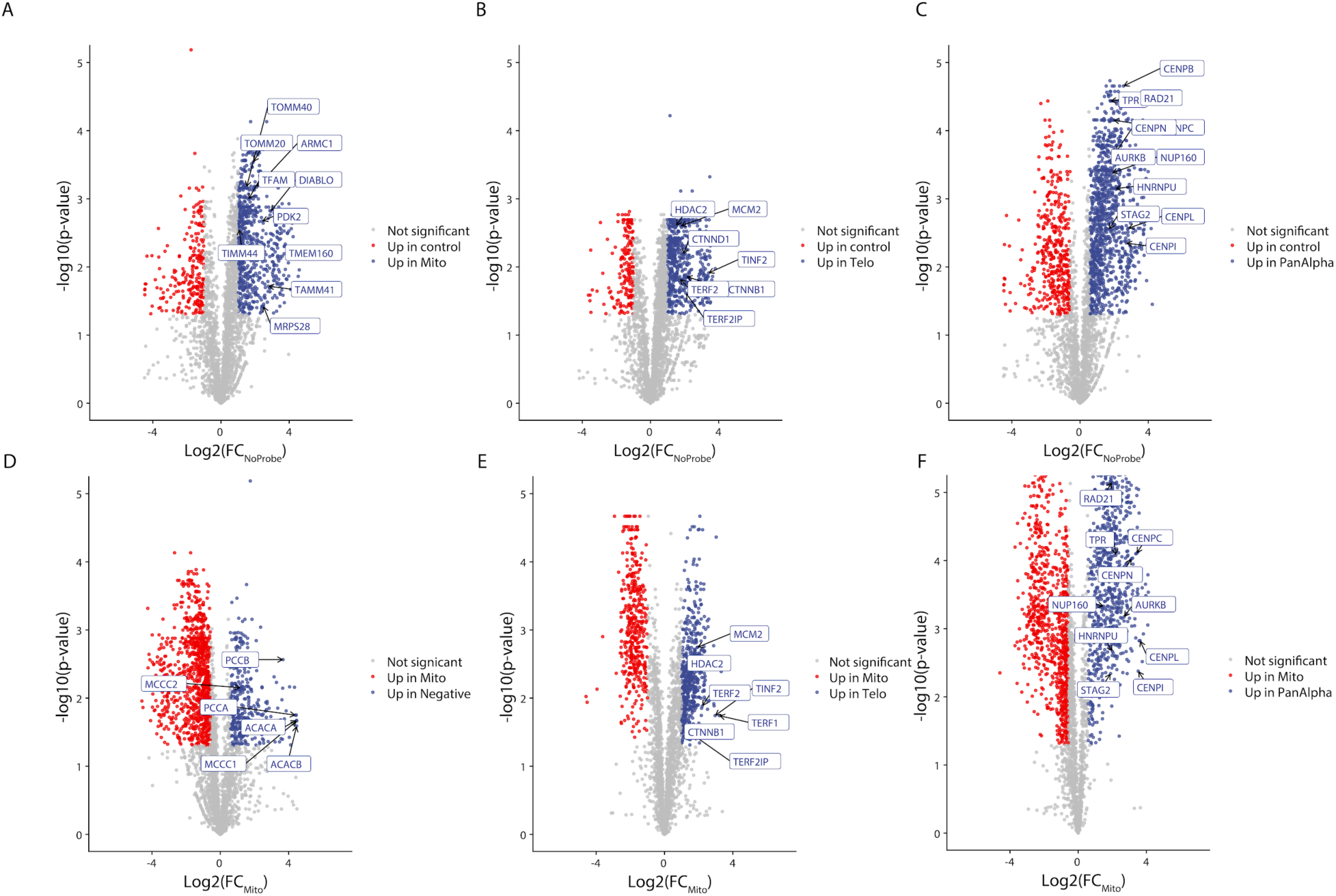
Relative quantitation for the multi-target DNA O-MAP proteomics experiment compared to no-probe control and mtDNA datasets. Volcano plots from multiplexed proteomics experiments with proteins of interest highlighted. A-C) Fold-changes and significance calculated compared to no probe. D-F) Fold-changes and significance calculated compared to mtDNA probe.

**Figure S5.**
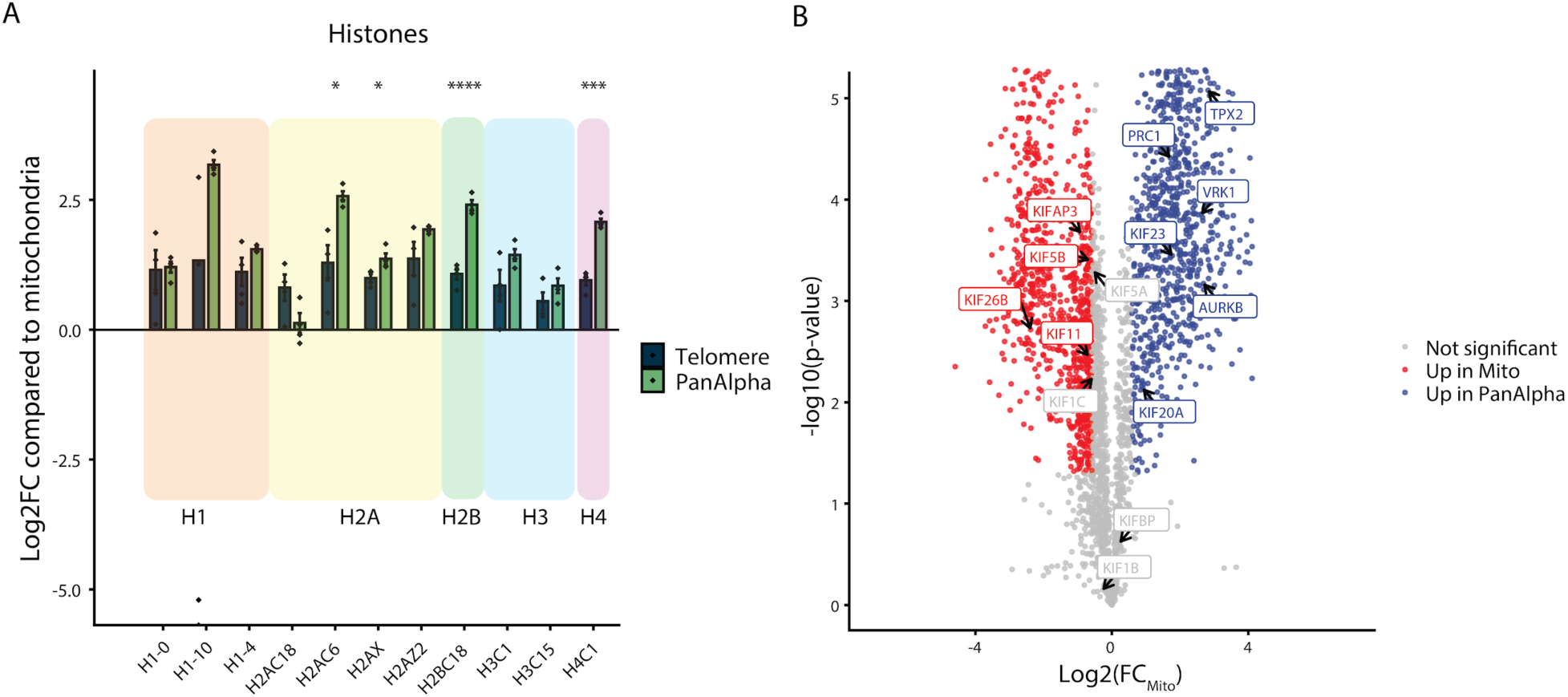
Comparison of histone proteins between telomere and pan-alpha probes. A) Log2 fold change of proteins compared to mitochondrial probe enriched histone complex proteins. Significance calculated based on Welch’s t-test for pairwise comparisons (p-value: *<0.05, **<0.01, ***<0.001, ****<0.0001). B) Volcano plot comparing the fold change of pan-alpha to the mtDNA probe with spindle proteins highlighted.

**Figure S6.**
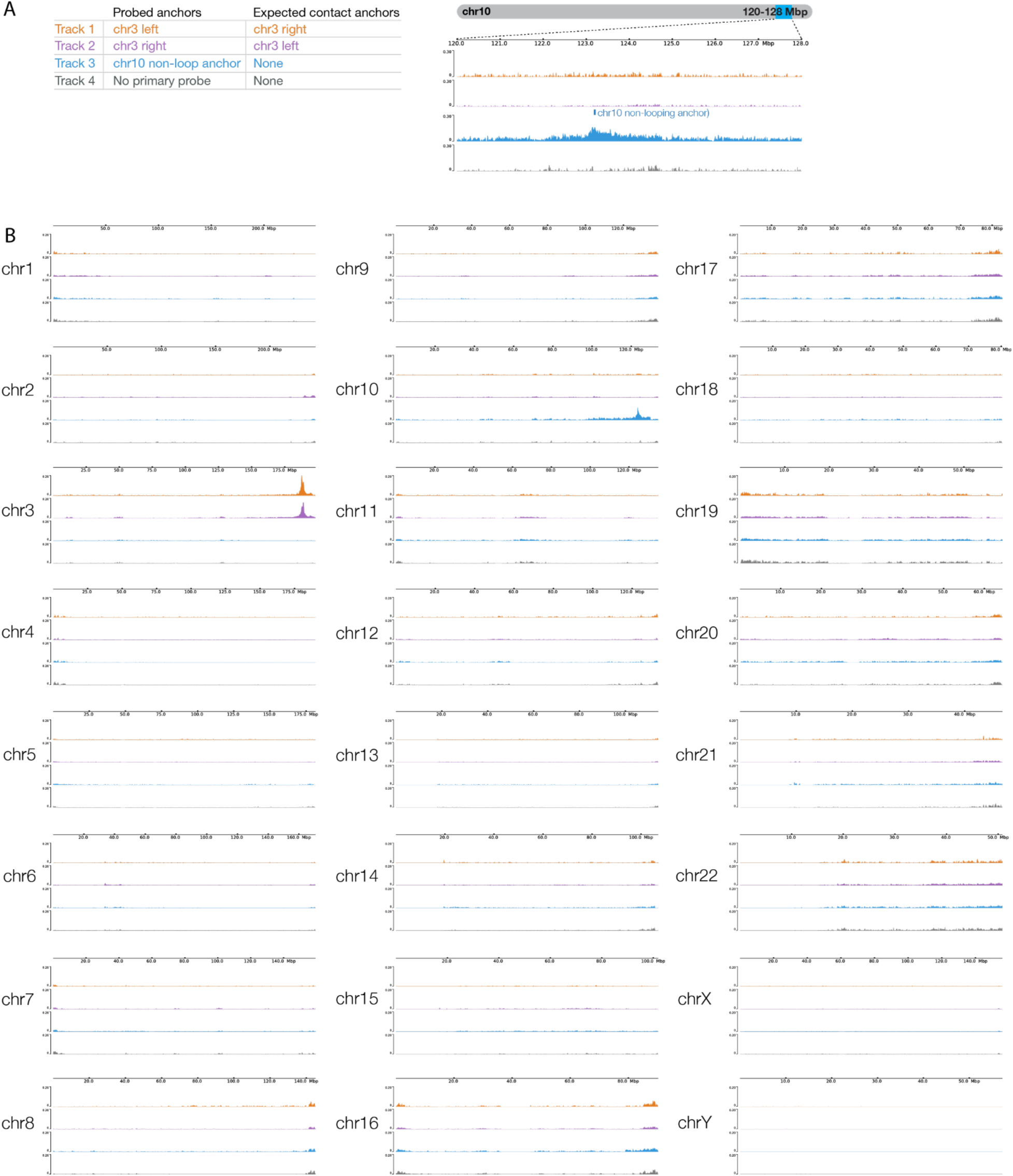
DNA O-MAP biotin purification sequencing of chr3 left, chr3 right, chr10 non-loop anchors, and no-primary-probe control. A) Table listing the three anchors (Track 1-3) and no-primary-probe control (Track 4) biotinylated by DNA O-MAP and their expected contact anchors (left). Biotin purification sequencing signals across the 8-Mb region on chromosome 10 corresponding to the chr10 non-loop anchor targeted (right). B) Biotin purification sequencing signals across every chromosome in the genome for this experiment.

**Figure S7.**
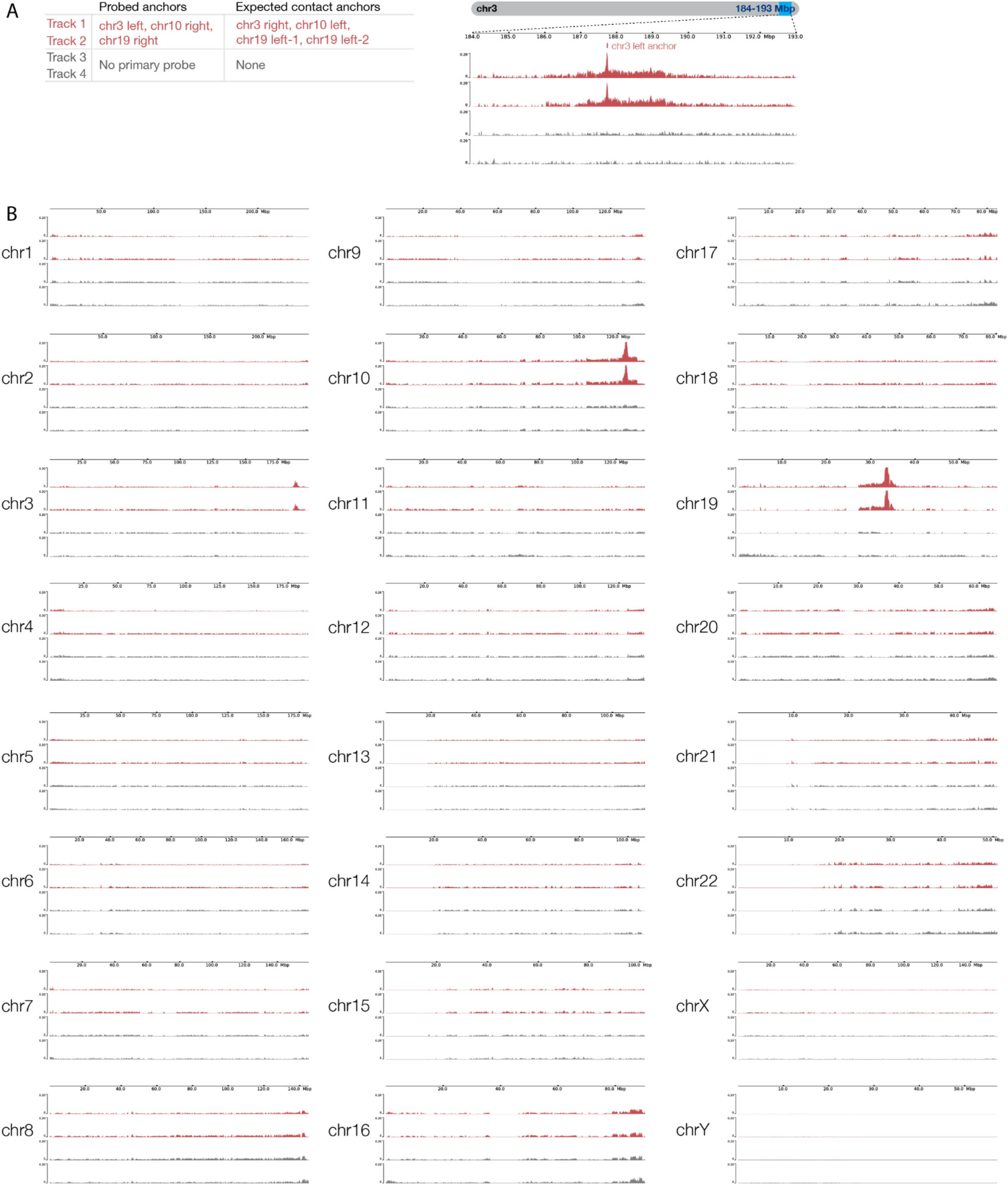
DNA O-MAP biotin purification sequencing of multiplexed targeting of chr3 left, chr10 right, chr19 right anchors, and no-primary-probe control in duplicates. A) Table listing the three anchors (Track 1-2) and no-primary-probe control (Track 3-4) biotinylated by DNA O-MAP and their expected contact anchors (left). Biotin purification sequencing signals across the 9-Mb region on chromosome 3 corresponding chr3 left anchor targeted in Track 1-2 (right). B) Biotin purification sequencing signals across every chromosome in the genome for this experiment.

**Figure S8.**
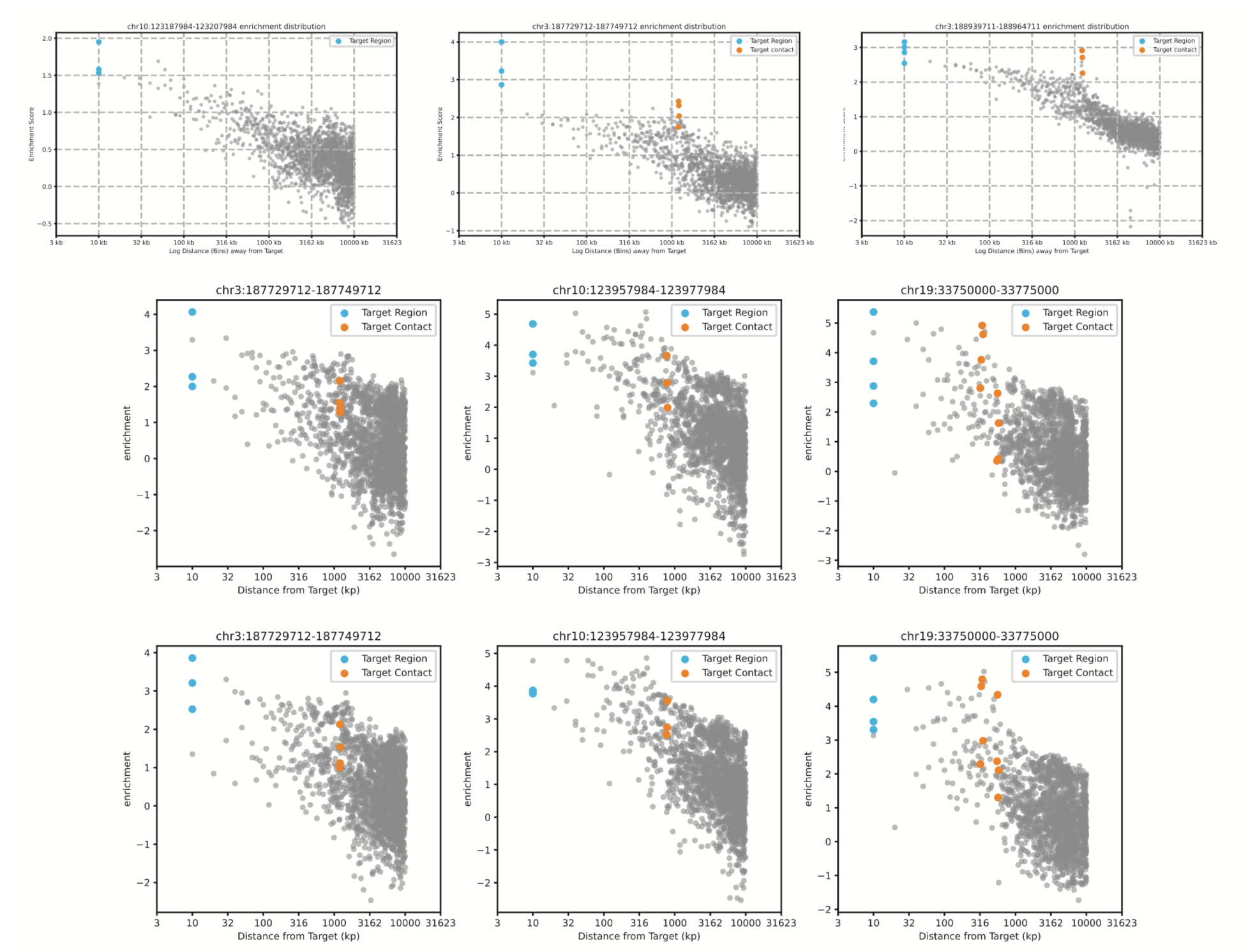
DNA O-MAP biotin purification sequencing enrichment score distribution of multiplexed targeting of chr3 left, chr10 right, chr19 right anchors. The scatter plots illustrate the enrichment score distribution for experiments targeting specific regions of chromosome 3, 10, and 19. The x-axes show the log distance from the target in kilobase pairs (kp), and the y-axes show the enrichment scores. Data points highlighted in blue represent the primary target region and the data points highlighted in orange represent the contact regions.

**Figure S9.**
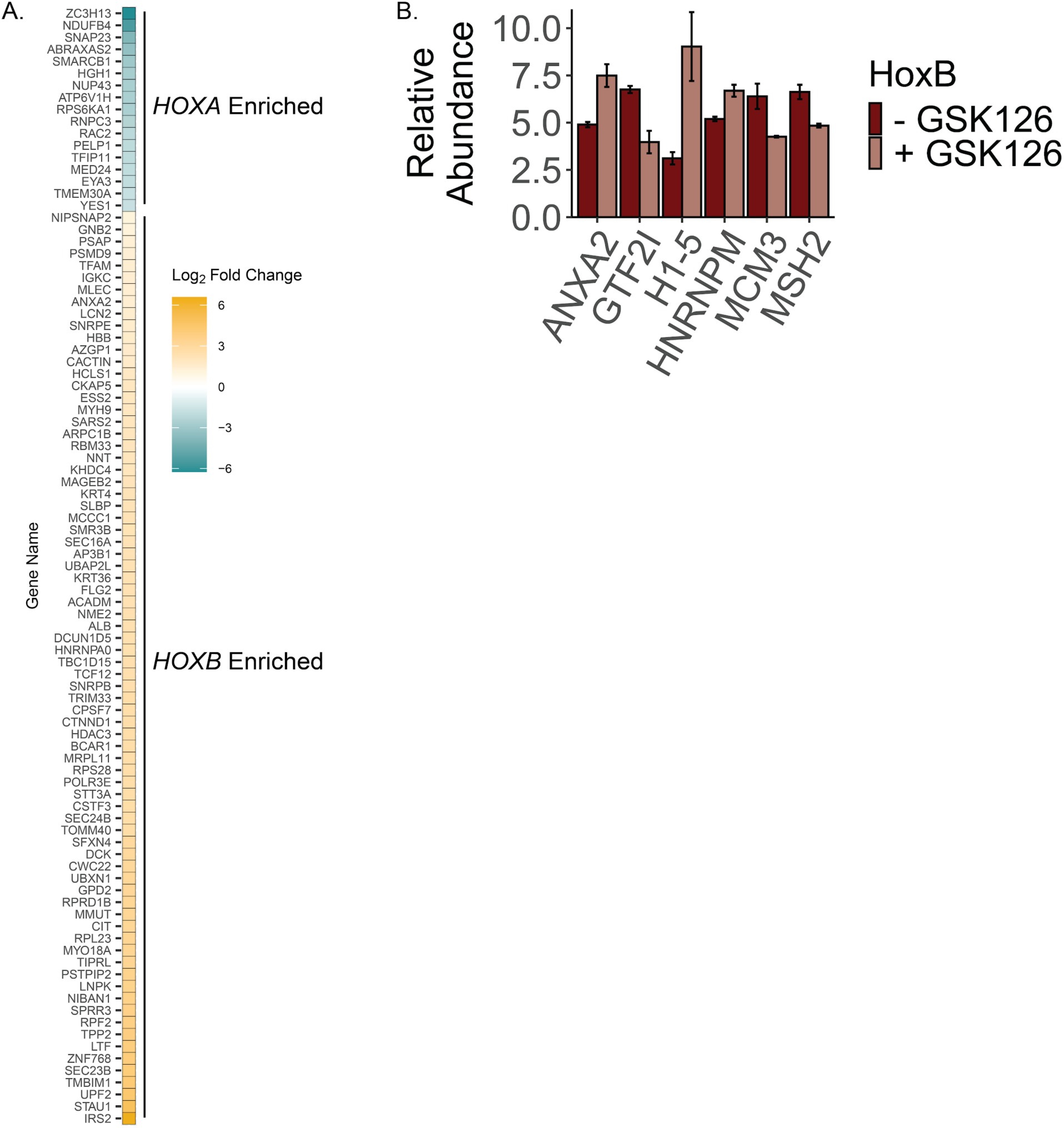
DNA O-MAP elucidates differences between the *HOXA* and HOX*B* proximal proteomes and changes to their proteomes following inhibition of EZH2 with GSK126. A) Heat map showing the Log2 fold change of all proteins that were scored as significantly enriched when comparing *HOXA* and *HOXB* proximal proteomes in K562 cells by DNA-OMAP. Adjusted p-value <0.05. B) Bar chart showing proteins with significantly altered abundance at *HOXB*, unique from *HOXA*, following treatment with GSK126 (Welch’s t-test, corrected p-value < 0.05).

**Figure S10.**
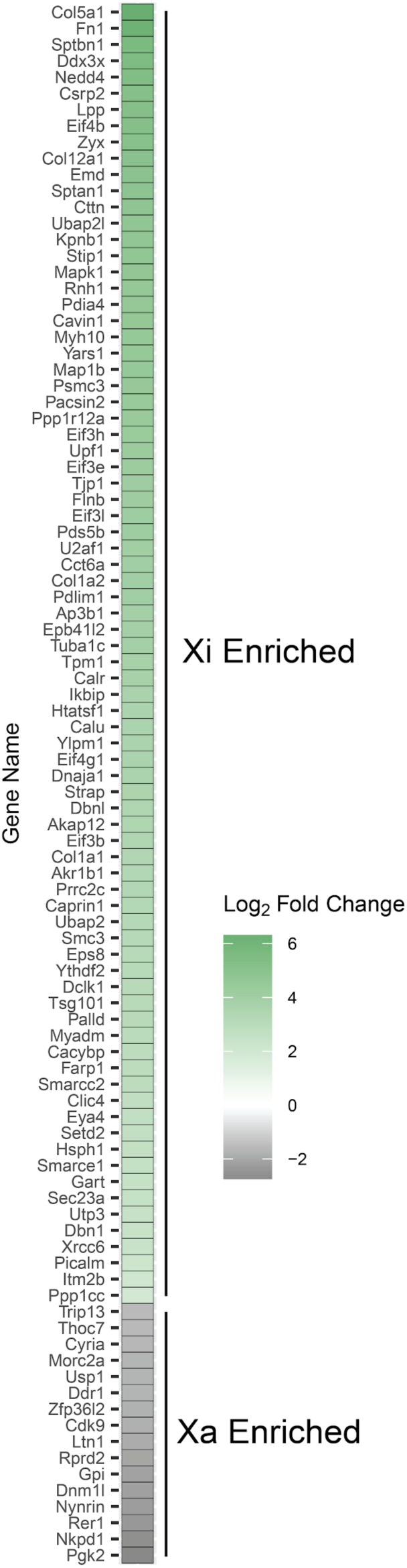
DNA O-MAP elucidates the differential proteomes of the X chromosome at homolog resolution. Heat map showing the Log2 fold change of all proteins that were scored as significantly enriched when comparing the active and inactive X chromosome homolog proximal proteomes in EY.T4 cells by DNA-OMAP. Adjusted p-value < 0.1.

**Supplementary Table 1.** Oligonucleotide probe sequences used in this work.

**Supplementary Table 2.** Proteomic data for the enrichment of telomere probe associated proteins.

**Supplementary Table 3.** Quantitative proteomic data for the multi-target DNA O-MAP proteomics experiment.

**Supplementary Table 4.** Summary of oligonucleotide probe target information.

**Supplementary Table 5.** Proteomic data for the differential enrichment of *HOXA* and *HOXB* associated proteins.

**Supplementary Table 6.** Quantitative proteomic data for the GSK126 treated *HOXA* proteomics experiment.

**Supplementary Table 7.** Quantitative proteomic data for the GSK126 treated *HOXB* proteomics experiment.

**Supplementary Table 8.** Proteomic data for the differential enrichment of Xi and Xa associated proteins.

## Notes

### Summary of Updates

We have added SI tables and made minor text revisions.

